# GABAergic neurons from the ventral tegmental area represent and regulate force vectors

**DOI:** 10.1101/2024.12.07.627361

**Authors:** Qiaochu Jiang, Konstantin I. Bakhurin, Ryan N. Hughes, Bryan Lu, Shaolin Ruan, Henry H. Yin

**Affiliations:** Department of Psychology and Neuroscience, Duke University, Durham, NC, 27708, USA; Department of Neurobiology, Duke University School of Medicine, Durham, NC, 27708, USA

**Keywords:** reward, ventral tegmental area, Pavlovian conditioning, force, GABAergic neurons, movement

## Abstract

The ventral tegmental area (VTA), a midbrain region associated with motivated behaviors, consists predominantly of dopaminergic (DA) neurons and GABAergic (GABA) neurons. Previous work has suggested that VTA GABA neurons provide a reward prediction, which is used in computing a reward prediction error. In this study, using in vivo electrophysiology and continuous quantification of force exertion in head-fixed mice, we discovered distinct populations of VTA GABA neurons that exhibited precise force tuning independently of learning, reward prediction, and outcome valence. Their activity usually preceded force exertion, and selective optogenetic manipulations of these neurons systematically modulated force exertion without influencing reward prediction. Together, these findings show that VTA GABA neurons continuously regulate force vectors during motivated behavior.

## Main

The ventral tegmental area (VTA) is a midbrain region that has traditionally been implicated in reward and motivation. Most VTA neurons are dopaminergic (DA) or GABAergic (GABA) (Nair-Roberts et al., 2008). The more common DA neurons show low tonic activity and occasional burst firing (Bunney et al., 1991; Guarraci & Kapp, 1999; Ungless & Grace, 2012), whereas the GABA neurons are tonically active with high firing rates and short-duration action potentials (Chieng et al., 2011; Lee et al., 2001; Steffensen et al., 1998). VTA GABA neurons represent a major output from the limbic basal ganglia (Breton et al., 2019; Yin, 2023). So far, most studies have focused on VTA DA neurons, and less is known about the contribution of VTA GABA neurons.

VTA GABA neurons receive projections from many areas, such as the prefrontal cortex and nucleus accumbens, and in turn project to many areas, including the prefrontal cortex, ventral pallidum, and habenula (Al-Hasani et al., 2021; An et al., 2021; Lammel et al., 2012; Margolis et al., 2012; Morales & Margolis, 2017; Sesack & Carr, 2002; Taylor et al., 2014; W.-L. Zhou et al., 2022). They often send collaterals to neighboring DA neurons, suggesting that they are in a position to influence dopaminergic signaling (Eshel et al., 2015; Matsui et al., 2014; Omelchenko & Sesack, 2009; Oster et al., 2015, 2015; Polter et al., 2018; Root et al., 2020). VTA GABA neurons have been implicated in both reward, aversion, locomotion, and sleep (Bouarab et al., 2019; Lee et al., 2001; Leemburg et al., 2018; Liu et al., 2012; Lowes et al., 2021; Puryear et al., 2010; Ranaldi, 2014; Yu et al., 2019, 2021). According to one hypothesis, while VTA DA neurons encode reward prediction errors, VTA GABA neurons encode reward prediction. Thus, the inhibitory projection from VTA GABA to DA neurons may implement the subtraction operation for computing reward prediction error (Cohen et al., 2012; Eshel et al., 2015; Pan et al., 2013). When the reward is predicted, VTA GABA neurons inhibit nearby DA neurons to reduce the effective teaching signal.

Other studies have shown that VTA GABA neurons are also responsive to aversive stimuli such as foot shock, air puff, and predator-like looming stimuli (Cohen et al., 2012; Liu et al., 2012; Root et al., 2020; Tan et al., 2012; Z. Zhou et al., 2019). Excitation of VTA GABA neurons could produce place aversion, and disrupt reward consumption and reward-seeking behavior (Shields et al., 2021; Tan et al., 2012; van Zessen et al., 2012). When continuous and precise behavioral measurements were used, many VTA GABA neurons were shown to represent kinematic variables such as pitch, yaw, and roll of the head, and play a causal role in steering the head during motivated behavior (Hughes et al., 2019). Optical excitation and inhibition of each population caused animals to move their heads in opposite directions. Such results suggest that these neurons can play a causal role in steering the head during motivated behavior.

Thus, previous work produced conflicting results on the contributions of VTA GABA neurons. A major limitation of previous studies is the lack of continuous behavioral measures. For example, studies investigating learning mechanisms often used appetitive Pavlovian conditioning tasks, yet the behavioral measurement in these studies was mainly limited to discrete events, such as licks in head-fixed mice or head entries and nose pokes in freely moving mice.

To elucidate the function of VTA GABA neurons in motivated behavior, we studied their activity in head-fixed mice in a stimulus-reward task similar to what was used in previous work on reward prediction (Hughes, Bakhurin, Petter, et al., 2020). Using load cells on the head-fixed setup, we measured continuous force exertion in mice while monitoring and manipulating neural activity using *in vivo* electrophysiology and optogenetics. We found that VTA GABA neurons represent and regulate force exertion. Although they are critical for modulating performance online, they are not needed for reward prediction or stimulus-reward learning.

## Results

On a head-fixation setup, we trained mice on a stimulus-reward task, in which a conditioned stimulus (200-millisecond tone -CS) is followed 1 second later by an unconditioned stimulus (US, 10% sucrose solution reward, Figure 1B). We measured the force exerted by the head of the mouse (K. I. Bakhurin et al., 2020; Hughes, Bakhurin, Barter, et al., 2020). Mice were not required to exert force to reach the waterspout; instead, force exertion reflected anticipatory and consummatory behaviors naturally generated during the task.

**Figure 1.**
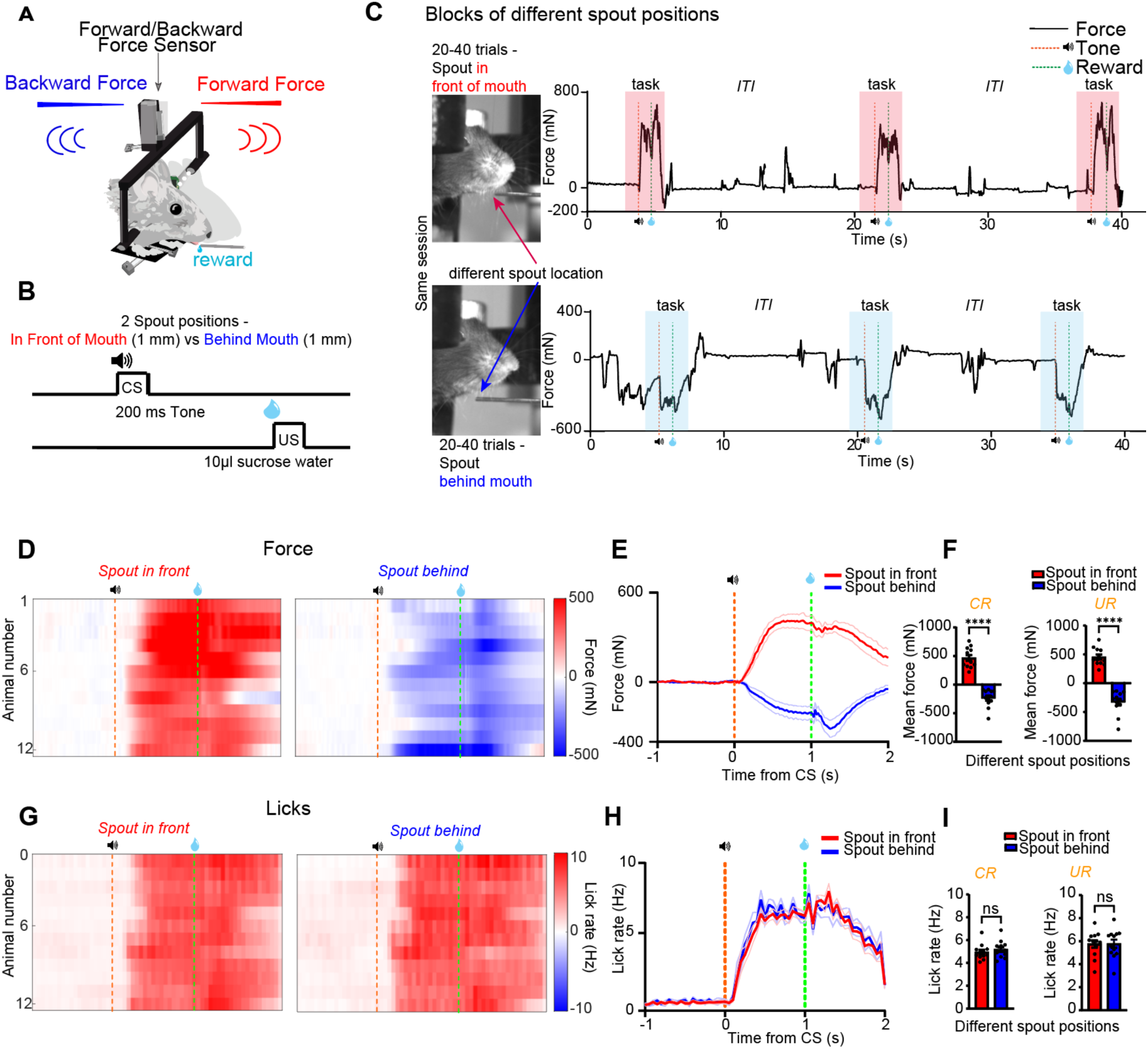
Changes in reward spout location induced force exertion in different directions without changing reward prediction. (A). Continuous measurement of forward and backward force exertion in mice restrained in a head-fixation device. (B-C) Pavlovian conditioning task design with different spout positions. (D-E) Bidirectional force exertion depends on the spout location within the same sessions. Mice demonstrated force exertion in direction aligned with the spout’s location (n = 12). (F) Mice exerted force in different directions as conditioned and unconditioned response (Left: CR, paired t-test, t = 9.473, p < 0.0001; Right: UR, paired t-test, t = 9.556, p < 0.0001). (G-H) Consistent licking behavior while spout location varied. (I) Left, same lick rate as the conditioned response (paired t-test, t = 1.758, p = 0.107). Right, the same lick rate as the unconditioned response (paired t-test, t = 0.0624, p = 0.951).

The spout for sucrose solution delivery was placed ∼1mm in front of or behind the lower jaw of the mouse, allowing it to drink from the spout easily (Figure 1C). Mice adjusted their force direction in response to the spout’s position: exerting forward force for front placements and backward force for rear placements. After training, mice consistently exerted force in response to the CS in the direction of the spout (Figure 1D), both for the conditioned response (CR; behavior after CS but before US) and the unconditioned response (UR; behavior after US; Figures 1D to 1F).

Despite varying force directions corresponding to reward spout placements, the mice showed similar licking rates in the tasks (Figures 1G and 1H). That is, manipulation of spout location altered force direction but not licking rate, the conventional measure of the CR on stimulus-reward tasks that is used to indicate reward prediction (Figure 1I).

### Distinct populations of VTA GABA neurons contribute to force exertion but not reward prediction

We chronically implanted drivable 16-channel optrodes into the VTA and recorded single-unit activity (VGAT-Cre mice, n = 8, DAT-Ai 32 mice, n = 4, Figure 2A). To confirm cell type identification, we optogenetically tagged some neurons using either a Cre-dependent excitatory opsin (channelrhodopsin, ChR2) to excite neurons or an inhibitory opsin (Cre-dependent anion-conducting channelrhosopin, stGtACR2) to inhibit neurons (n = 351 total neurons; 24 stGtACR2-tagged GABA neurons and 17 ChR2-tagged GABA neurons; Figure S2) in VGAT-Cre mice. The optogenetically tagged neurons have similar waveforms as other neurons classified as putative GABA neurons, showing short-duration action potentials and high tonic firing rates (Figures S2B and S2I). Putative GABA neurons (n = 178) showed a firing rate of 18.27 ± 1.03 Hz (mean ± sem) with narrow spike waveforms (full width at half maximum (FWHM): 142 ± 4.5 ms), and unclassified neurons (n = 173) showed a firing rate of 5 ± 0.32 Hz with wider spike waveforms (FWHM: 239 ± 12.7 ms; Figure S1).

**Figure 2.**
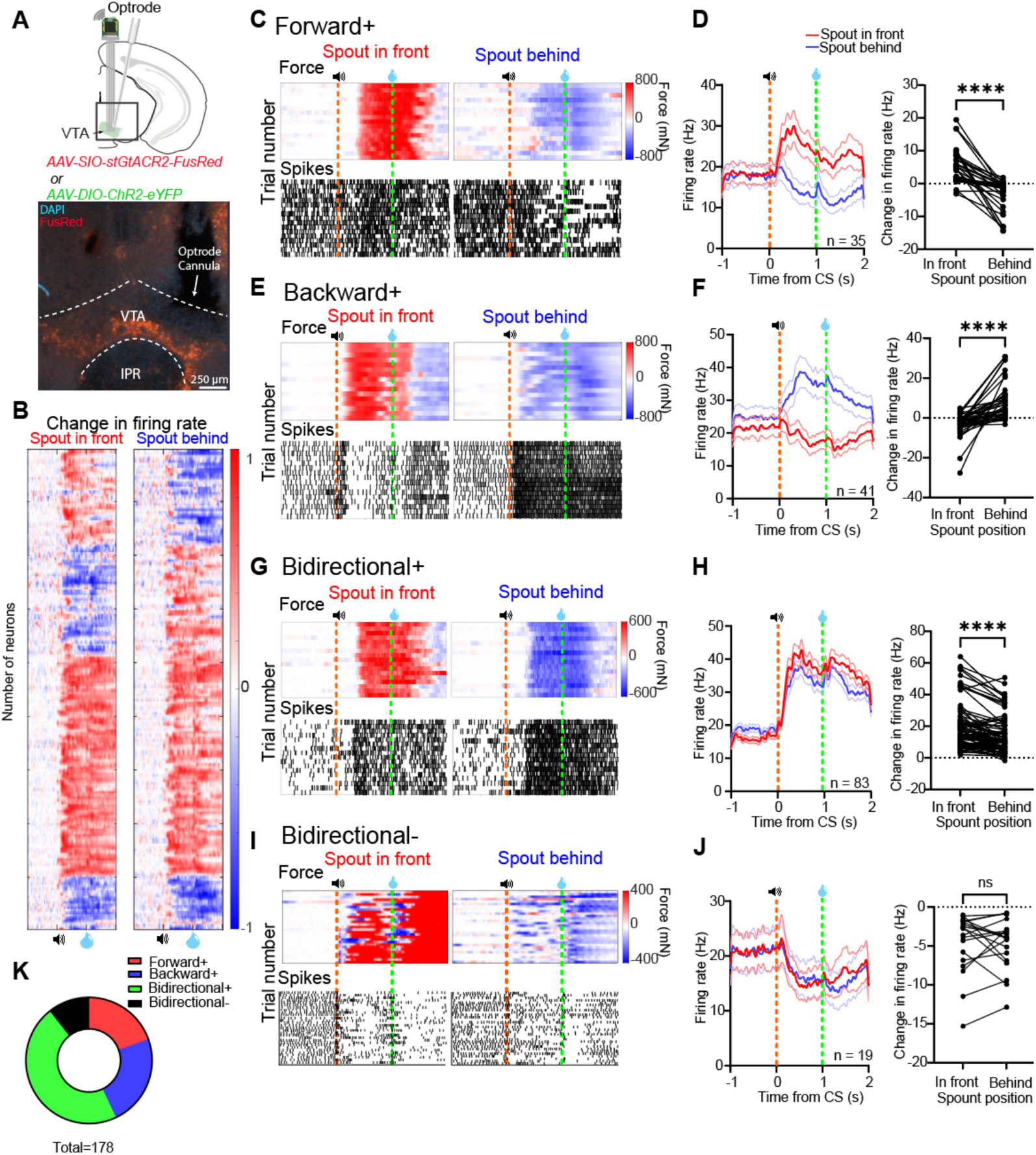
Distinct force tuning profiles of VTA GABA neurons. **(A)** *Top*, *in vivo* recording from GABAergic neurons by implanting an optrodes above VTA. *Bottom*, expression of AAV-SIO-stGtACR2-FusRed in VGAT+ neurons. **(B)** Activity profiles of GABAergic neurons showing normalized changes in firing rates in Pavlovian conditioning with spout location variation. Red, increase from baseline; blue, decrease from baseline. Baseline is defined as the mean FR during 1 second before CS (*Left*: in front spout placements; *Right*: behind spout placements; n = 178 neurons). **(C)** Representative example of a Forward+ neuron increasing firing rate to forward force exertion but not to backward force exertion. Each stick is a spike. **(D)** *Left*: Forward+ population increased firing rate when the mice exerted forward force.*Right*: Forward+ neurons showed higher firing rates when the spout was in front than when the spout was behind (paired t-test, t = 7.022, p < 0.0001; n = 35 neurons). **(E-F)** Representative example and population summary showed that Backward+ neurons showed an increased firing rate to backward force exertion but not to forward force (paired t-test, t = 6.139, p < 0.0001; n = 41 neurons). **(G-H)** Bidirectional+ population increased firing rates when the mice exerted forward and backward forces, with a more significant increase in firing rate in response to forward force (paired t-test, t = 6.815, p < 0.0001; n = 83 neurons). **(I-J)** Bidirectional-neurons decreased firing rates when the mice exerted forward and backward forces (paired t-test, t = 0.269, p = 0.791; n = 19 neurons).

We examined the relationship between the activity of putative GABA neurons and the measured force from load cell sensors. Using an unbiased hierarchical clustering algorithm, we observed four populations of VTA GABA neurons based on their distinct force-tuning profiles (Figures 2B and S1). The firing rates of all four populations were closely related to measured force, but they exhibited distinct tuning based on the direction of force exertion. One population (Forward+ neurons, n = 35) increased firing for forward force generation (when the spout was placed in front) and decreased firing for backward force generation (when the spout was placed behind; Figures 2C, 2D, and S6A). The second population (Backward+ neurons, n = 41) showed the opposite pattern: their firing rate increased during backward force exertion and decreased during forward force (Figures 2E, 2F, and S6B). These two populations were tuned for continuous force exertion but had opposite direction preferences (Figures 3B to 3E). The third population (Bidirectional+ neurons, n = 83) increased firing regardless of force direction (Figures 2G, 2H, and S6C). The fourth population (Bidirectional-neurons, n = 19) decreased firing when the mouse exerted forward and backward forces (Figures 2I, 2J, and S6D). Bidirectional+ and Bidirectional-neurons did not show direction preference, but showed activity that is highly correlated with the absolute value of force exertion in either forward or backward direction (Figures 3H, 3L, S6C, and S6D). Representative and tagged examples of different types of VTA GABA neurons are shown in Figures 3B, 3D, 3F, 3J, and S3.

**Figure 3.**
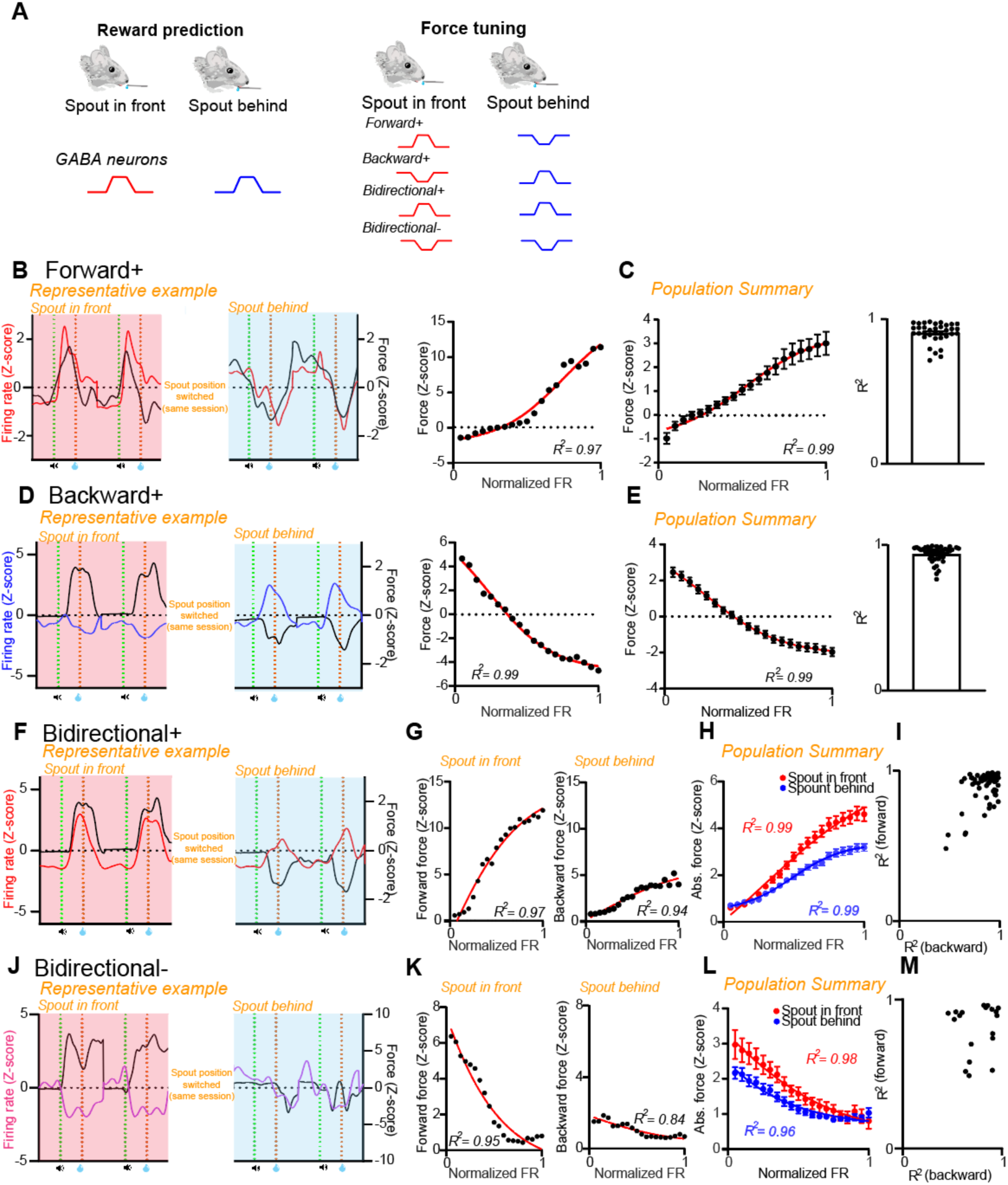
GABAergic neurons represent force continuously independent of reward prediction. **(A)** Schematic illustrations of reward prediction hypothesis and force tuning hypothesis of GABAergic neurons in different spout placement conditions. **(B)** *Left*: a representative Forward+ neuron activity (red) closely represented forward force exertion (black) during tasks. *Right*: the example neuron’s firing rate was highly tuned for forward force exertion (R^2^ = 0.97). **(C)** *Left*: Forward+ population was positively tuned for forward force amplitude in the task (R^2^ = 0.99, n = 35). *Right*: individual Forward+ were robustly tuned for forward force exertion. **(D-E)** Backward+ example neuron **(D)** and population **(E)** were tuned for backward force exertion (R^2^ = 0.99, n = 41). **(F-I)** Bidirectional+ example neuron (**F** & **G**) and population (**H** & **I**) showed positive tuning with the absolute value of forward and backward force exertion (forward force: R^2^ = 0.99, n = 83; backward force: R^2^ = 0.99, n = 83). **(J-M)** Bidirectional-example neuron **(J & K)** and population **(L & M)** exhibited a negative tuning for the absolute value of forward and backward force exertion (forward force: R^2^ = 0.98, n = 19; backward force: R^2^ = 0.96, n = 19)

Our spout placement manipulations therefore induced mice to exert force in opposite directions while the licking rate remained consistent. Although mice showed consistent reward prediction, their performance varied in the direction of force exertion. This feature is also captured by the electrophysiology recording data, which reveal distinct activity patterns of different GABA neuron types depending on the direction of force exertion (Figure S4).

To see whether the GABA neurons representing force may play a causal role in generating the force, we examined the temporal relationship between neural activity and force generation. Forward+ neurons and Backward+ neurons usually led force exertion (Figures S5A and S5B). Bidirectional+ neurons also led force generation (Figure S5C), but the activity of Bidirectional-neurons lagged force generation (Figure S5D).

### VTA GABA neurons maintain force direction preferences in spontaneous movements

Despite being head-fixed, the mice did not sit still during the intertrial intervals (ITI). Occasionally, they tried to move, as indicated by force measurements. We characterized such movements without licking during the intertrial interval as “spontaneous movements” (Figure 4A). These movements were usually characterized by smaller force and shorter duration compared to movements during the trial (Figure 4B). If VTA GABA neurons represent force exertion variables, they should maintain their force tuning properties when generating spontaneous movements. This hypothesis was confirmed: the relationship between neural activity and force was similar regardless of whether the movements were spontaneously emitted or in response to cues and rewards (CR and UR; Figure 4).

**Figure 4.**
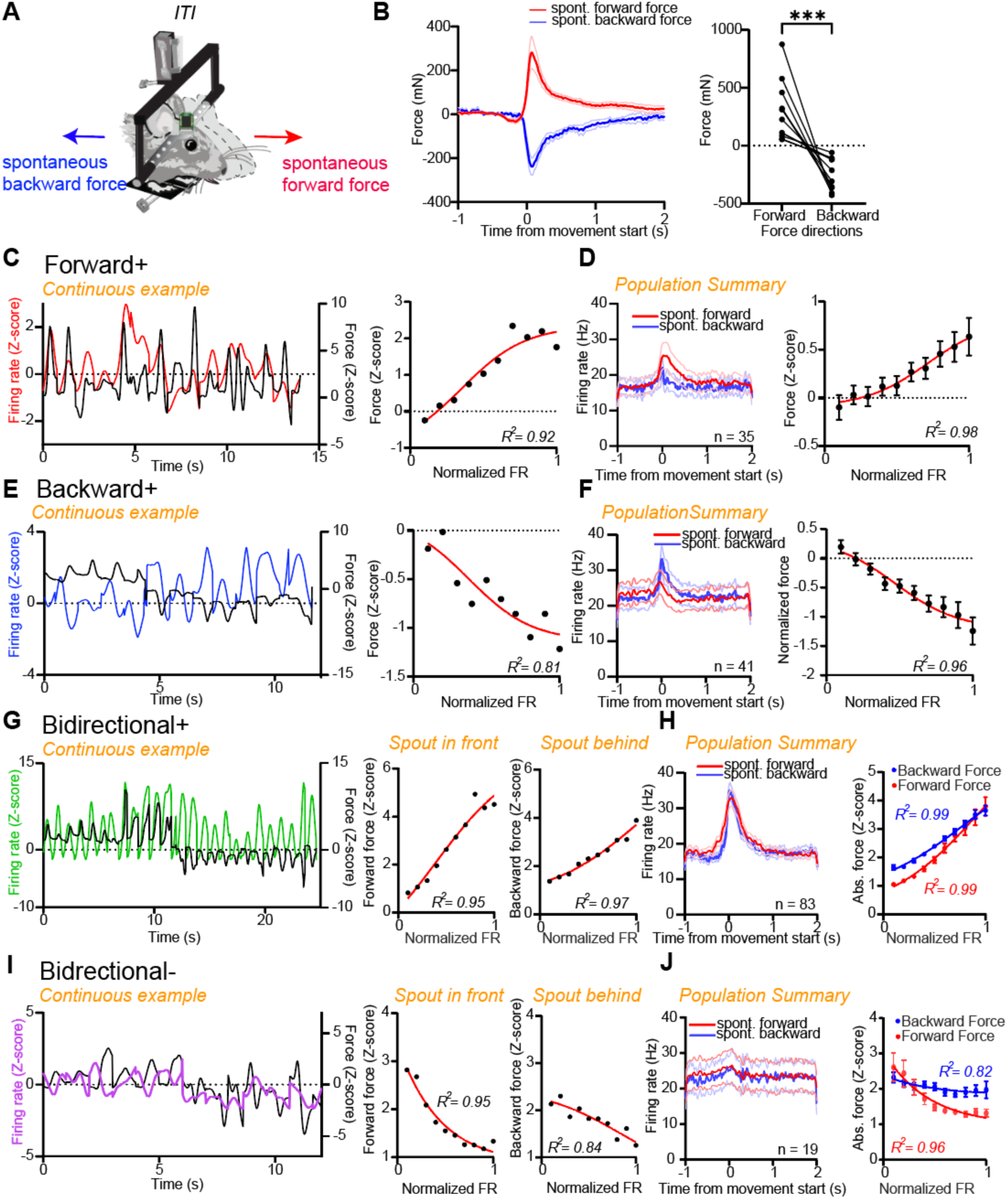
Similar force tuning across various GABA neuron populations during spontaneous movements. (A) Schematic illustration of mice moving spontaneously and exerting forward and backward forces outside of the task intervals. (B) Left: mice spontaneously exerted forward and backward forces. Right, average amplitude of spontaneous forward and backward force of individual mice (n = 12). (C) Left: the example Forward+ neuron’s firing activity followed spontaneous forward force exertion. Right: the example neuron showed a robust tuning for spontaneous forward force exertion (R2 = 0.92). (D) Left: Forward+ population activity to spontaneous forward and backward forces. Right: Forward+ population was tuned for spontaneous forward force (R2 = 0.98, n = 35). (E-F) Backward+ example neuron (R2 = 0.81) and population were tuned for spontaneous backward force (R2 = 0.96, n = 41). (G-H) Bidirectional+ example neuron and population were tuned for both spontaneous forward and backward forces (forward: R2 = 0.99; backward: R2 = 0.99, n = 83). (I-J) Bidirectional-example neuron and population exhibited negative tuning for absolute values of spontaneous forward and backward forces (forward: R2 = 0.96; backward: R2 = 0.82, n = 83).

### Change in GABA activity to aversive stimuli is explained by force tuning

We also tested whether the activity of VTA GABA neurons depends on outcome valence. Previous studies found that VTA GABA neurons change their firing rates in response to aversive stimuli (Cohen et al., 2012; Root et al., 2020; Tan et al., 2012). We also tested the response of VTA GABA neurons to aversive stimuli (delivery of air puffs) while measuring force exertion (Figure 5A). We observed that unpredicted air puffs generated consistent backward movements away from the stimulus, followed by a rebound forward movement (Figures 5B to 5D). Indeed individual neurons maintained their force tuning in response to air puffs (Figures 5E to 5L and Figure S7). Thus, the activity of VTA GABA neurons are independent of motivational context or outcome valence (reward or aversion).

**Figure 5.**
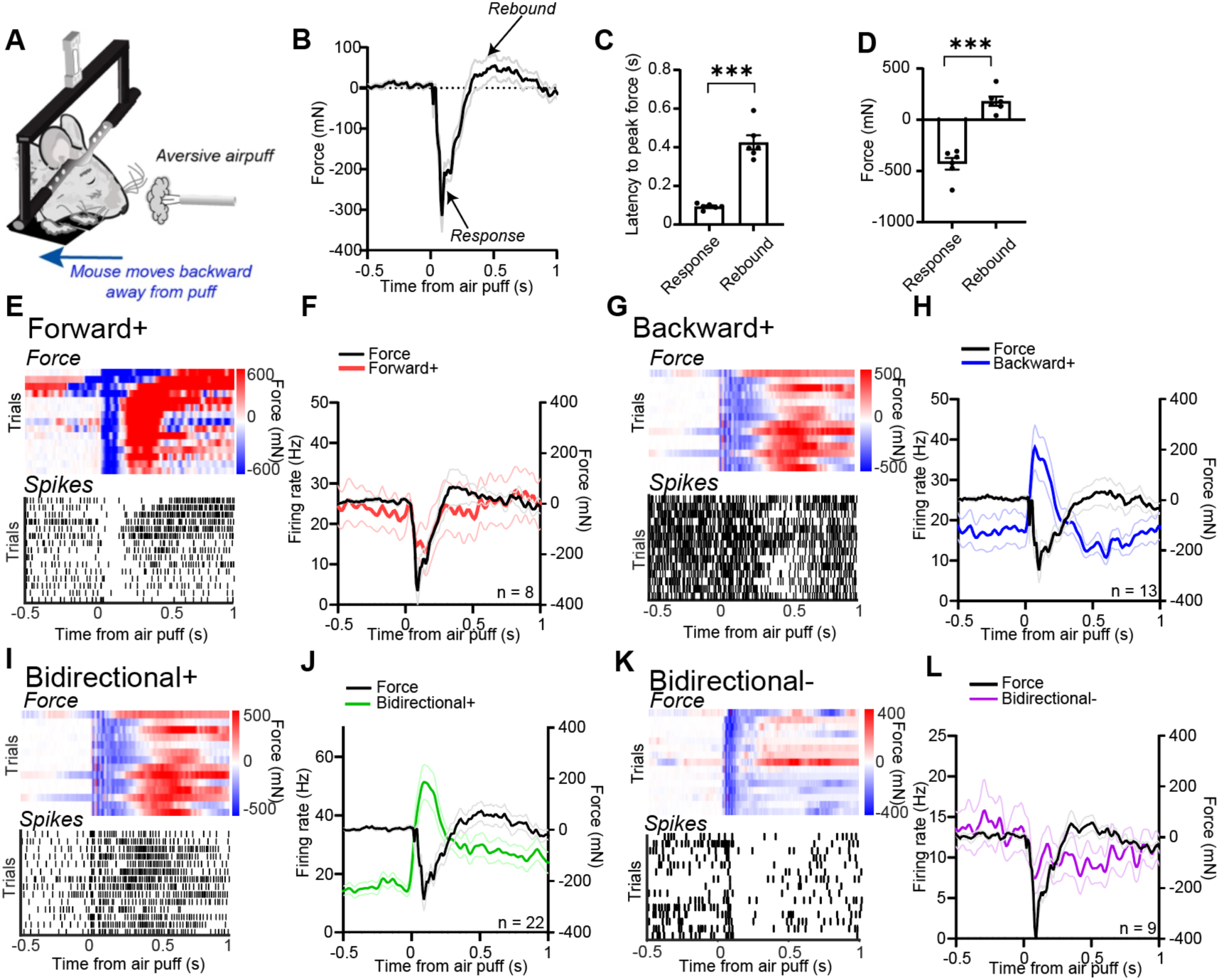
GABA neurons maintain force tuning in response to aversive air puffs. (A) Schematic of mice subjected to mild, unpredicted air puff to the face in sessions independent of Pavlovian conditioning. (B) Summary of force exertion across all mice (n = 6 mice). (C) Sequence of force exertion in mice, depicting initial backward followed by forward force in response to air puffs (paired t-test, p = 0.0002, t = 10.01, n = 6). (D) Mice exerted backward force as response and forward force as rebound (paired t-test, p = 0.0004, t = 8.28, n = 6). (E) *Top*: a representative example session of force as a response to air puff. *Bottom*: a representative Forward+ neuron responding to the air puff. Each tick is a spike. (F) Forward+ population activity in response to the air puffs (n = 10 neurons). (G-L) Same tradition as (E & F). Example and population activity of Backward+ (G & H), Bidirectional+ (I & J), and Bidirectional-(K & L) neurons in response to the air puff.

### Change in GABA activity in response to omission of predicted reward can be explained by force tuning

In well-trained mice, we omitted the reward (US) on 20% to 50% of the trials, while the spout was consistently placed in front of the mice (Figure S8A). We compared force and neural activity from rewarded and unrewarded trials. Mice abruptly stopped force exertion when the reward was omitted (Figure S8B). Forward+ and Bidirectional+ neurons did not show elevated firing rate during reward omission as they did when the mice consumed the reward (Figures S8C and S8E). Backward+ and Bidirectional-neurons did not significantly alter their activity (Figures S8D and S8F). These findings demonstrate that the change in force exertion to omitted rewards could be explained by the activity of force-tuned VTA GABA neurons.

### Optogenetic excitation of VTA GABAergic neurons suppressed anticipatory forward force exertion

Our electrophysiological results showed that the activity of VTA GABA neurons represented distinct components of the force vector and usually preceded force exertion. They suggest that these neurons may play a causal role in modulating performance online. To test this hypothesis, we used optogenetics to stimulate VTA GABA neurons during behavior. We injected a Cre-dependent excitatory opsin (ChR2, n = 6) or GFP (n = 4) into the VTA of VGAT-Cre mice (Figures 6A and S10A). This strategy allowed us to excite GABA(VGAT+) neurons selectively. We delivered light stimulation (40Hz, 40 Pulse, 15ms) at different time points. On 20% of trials, we delivered a 1-second stimulation during the CS-US trace interval (Figure 6B). On 20% of trials, we delivered a 1-second stimulation immediately after reward delivery (Figure 6G).

**Figure 6.**
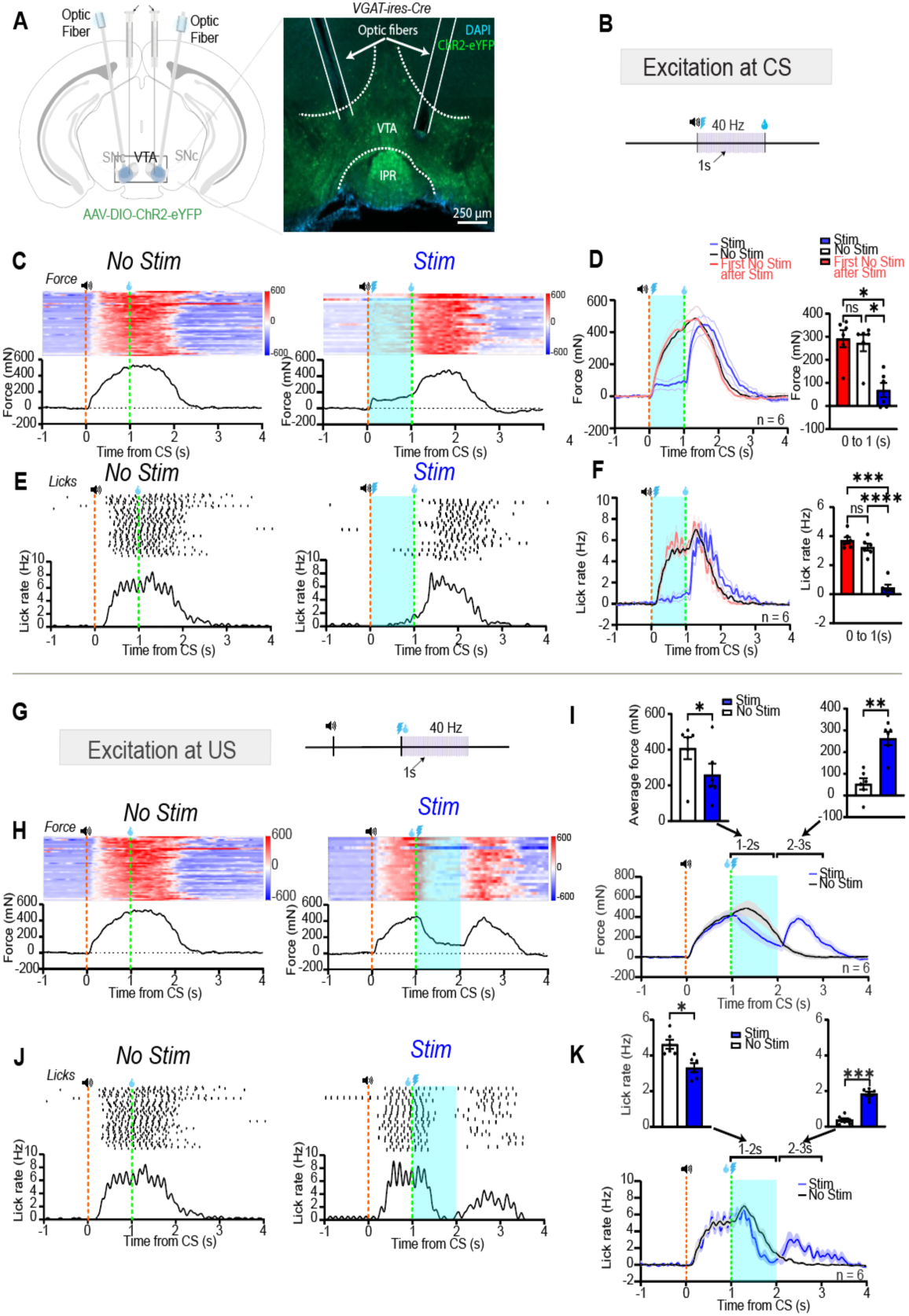
Optogenetic activation of GABAergic neurons modulates force exertion. (A) Left: schematic illustration of virus injection and optic fibers implanted into bilateral VTA. Right: histological verification of the optic fiber tip placement into VTA. (B) Schematic illustration of delivering stimulation between the CS-US interval. (C) Force exertion of a representative mouse in trials without stimulation (Left) and with stimulation (Right). (D) Left: mice force exertion in different trial types. Right: a one-way ANOVA showed mice exerted different force amplitude in different trials (F (1.032, 5.162)21.19, p = 0.0053). Post-hoc analyses showed that mice exerted less force during stimulation (paired t-test, t = 4.788, p = 0.0049, n = 6 mice); mice exerted similar force for trials without stimulation and first trials without stimulation following stimulation at CS (paired t-test, t = 1.961, p = 0.1071, n = 6 mice). (E) Lick rate of a representative mouse in trials without stimulation (Left) and with stimulation (Right). (F) Left: mice licking behavior in different trial types. Right: mice licked less during stimulation (paired t-test, t = 21.97, p < 0.001, n = 6); mice similarly in trials without stimulation and first trials without stimulation following stimulation at CS (paired t-test, t = 2.378, p = 0.0633, n = 6 mice). (G) Schematic illustration of delivering stimulation at the time of the US. (H) Force exertion of a representative mouse in trials without stimulation (Left) and with stimulation at the US (Right). (I) Mice’s force was suppressed during stimulation (paired t-test, t = 3.006, p = 0.0299, n = 6 mice) but rebounded after the cessation of stimulation (paired t-test, t = 5.354, p = 0.0031, n = 6 mice). (J) Licking of a representative mouse in trials without stimulation (Left) and with stimulation at the US (Right). (K) Licking was suppressed during stimulation (paired t-test, t = 2.819, p = 0.0371, n = 6 mice) but rebounded after the cessation of stimulation (paired t-test, t = 10.52, p = 0.0001, n = 6 mice).

Optogenetic stimulation during the CS-US interval immediately suppressed anticipatory forward force exertion as well as anticipatory licking (Figures 6C to 6F; Supplementary Video 1). This suppression was confined to the stimulation period. Force exertion and licking resumed as soon as stimulation stopped. Stimulation had no effects in GFP-expressing control mice (Figures S10B and S10C).

Stimulation did not have a long-term effect. Stimulation during the CS-US interval had no impact on performance during the subsequent no-stimulation trial (Figures 6D and 6F). This finding does not support the idea that VTA GABA neurons signal reward prediction (Cohen et al., 2012; Eshel et al., 2015). Its impact is limited to performance modulation in real time rather than learning.

### Optogenetic excitation of VTA GABAergic neurons suppressed reward consumption

To test if optogenetic excitation of GABA neurons had a similar effect on consummatory behavior, we also delivered stimulation at the time of reward delivery (Figure 6G). Stimulation at reward reduced the forward force during reward consumption (Figure 6H and 6I; Supplemental Video. 2). There was also a decrease in licking during stimulation (Figures 6J and 6K; Supplemental Video. 2). Upon cessation of stimulation, however, there was a notable rebound in both forward force and licking (Figures 6I and 6L). Stimulation did not affect anticipatory force exertion or licking on the next no-stimulation trial (Figure S9). In control mice with no opsins, stimulation had no effect on behavior (GFP control, Figures S10D and S10E).

We also tested the impact of optogenetic stimulation in the absence of a reward (Figure 7A). In the same session, rewards were omitted at a 50% probability following the CS presentation. On reward omission trials, stimulation was also delivered with a 50% probability at the expected time of the reward. On these trials stimulation still effectively suppressed forward force exertion (Figures 7B and 7C). Following the termination of stimulation, there was a clear rebound in both force exertion (Figure 7C) and licking (Figure 7E) despite the absence of any reward delivery and consummatory licking. In other words, regardless of the motivational context, reward prediction, and reward consumption, all mice suppressed force exertion during stimulation. The behavior showed a “rebound” pattern once stimulation stopped. These effects were not observed in control mice (Figure S11).

**Figure 7.**
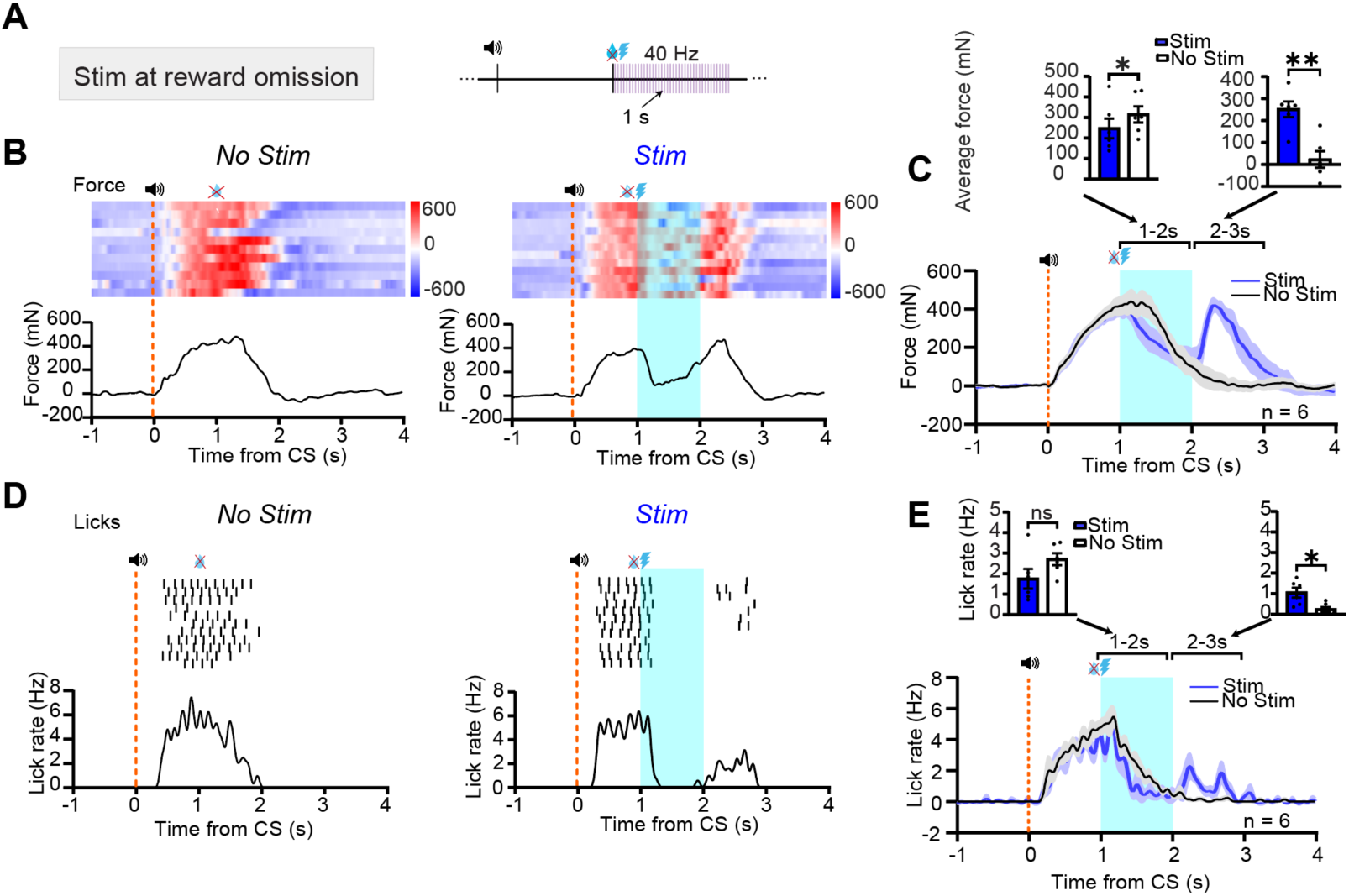
Stimulating VTA GABA neurons during reward omission experiments. **(A)** Schematic illustration of stimulation at reward omission. **(B)** Force exertion of a representative mouse in trials without stimulation (*Left*) and with stimulation (*Right*) when the US was omitted. **(C)** Mice exerted less force in trials with stimulation for 1 to 2 seconds after CS (paired t-test, t = 3.182, p = 0.0245, n = 6 mice) then showed a force rebound after stimulation (paired t-test, t = 4.554, p = 0.0061, n = 6 mice). **(D)** Licking of the representative mouse. **(E)** Mice licked similarly in trials with and without stimulation during 1 to 2 seconds after CS (paired t-test, t = 1.978, p = 0.1049, n = 6 mice) but licked more during 2 to 3 seconds after CS in trials with stimulation (paired t-test, t = 3.306, p = 0.0213, n = 6 mice).

## Discussion

Our results revealed four distinct populations of VTA GABA neurons with distinct force tuning properties. These neurons vary their firing rates during behavior depending on the direction of force exertion, even when reward prediction remains the same (Figures 1 and 2). Many of them (Forward+, Backward+) show strong direction preference, whereas others (Bidirectional+, Bidirectional-) show activity representing the absolute magnitude of force exertion (Figure S6).

VTA GABA neurons maintain their force tuning under different conditions, e.g., spontaneous behavior (Figure 4) and aversive air puff (Figure 5). In addition, when the rewards were omitted, 28 out of 29 forward force-tuned neurons showed a decrease in firing rate, accompanied by reduced force. Their activity is also independent of learning and reward prediction. Usually, their activity slightly leads the force variable, suggesting they are involved in force generation (Figure S5). Direct support for a causal role of VTA GABA neurons in force generation in this task was provided by our optogenetic experiments, which demonstrated that activation of VTA GABA neurons precisely modulated force exertion for both anticipatory and consummatory behaviors (Figures 6 and 7).

VTA DA neurons, which receive inhibitory projections from VTA GABA neurons, are often believed to signal reward prediction error, the difference between actual reward and reward prediction (Cohen et al., 2012; Schultz, 1998). Uchida and colleagues proposed that VTA GABA neurons, which inhibit DA neurons, encode reward prediction and play a role in computing reward prediction error (Cohen et al., 2012; Eshel et al., 2015). However, recent results have questioned this interpretation. Studies have found that VTA DA neurons signal salience and kinematics independent of learning or reward prediction (Barter et al., 2015; Berridge, 2007; Hughes, Bakhurin, Petter, et al., 2020; Kutlu et al., 2021, 2023; Root et al., 2020; Wang & Tsien, 2011). VTA GABA neurons also represent rotational kinematics (pitch, yaw, roll) in freely moving mice and play a causal role in steering the head (Hughes et al., 2019). These results suggest that VTA neurons are critical for performance rather than learning. In accord with such findings, our results demonstrate that, in head-fixed mice, VTA GABA neurons represent force vectors in performance without directly contributing to learning or reward prediction. Instead of encoding reward prediction, as previously argued, VTA GABA neurons continuously modulate performance in real time.

VTA GABA neurons are assumed to be projection neurons that send axons to distal structures, but we cannot rule out the possibility that some of them are local interneurons that mainly synapse on neighboring neurons within the VTA. The four populations of VTA GABA neurons we identified based on force tuning have similar spike waveforms but higher tonic firing rates compared to unclassified neurons (Figure S1). It is unknown whether they have distinct molecular signatures. In addition to GABA and DA neurons, the VTA also contains glutamatergic neurons as well as neurons that co-release glutamate and GABA(An et al., 2021; Ma et al., 2023; McGovern et al., 2021; Root et al., 2018, 2020; Yoo et al., 2016). It is also unclear whether the glutamatergic/ glutamate-GABA neurons would also exhibit force tuning.

Recent work has shown that VTA DA neurons also demonstrate force tuning, but their activity predicts a change in force rather than force per se (K. Bakhurin et al., 2023). It is possible that the DA neurons may modulate upstream structures like the ventral striatum, and the striatal output is integrated to yield a force command from the VTA GABA neurons. It is also possible that DA output could reflect the derivative of the GABA output. If the output of the VTA GABA neurons send descending commands to exert force in different directions, then their neighboring DA neurons, which receive direct GABAergic projections from them, may implement an adaptive gain mechanism by projecting back to upstream ventral striatal areas such as the nucleus accumbens.

Cohen et al. (2012) described VTA GABA neurons that appear to signal reward prediction. According to them, these “type II” neurons show sustained activation following the presentation of the CS, and their outputs can be used to compute reward prediction errors (e.g. actual reward minus predicted reward). They classified neurons with sustained inhibited activity in response to the CS as “type III” neurons.

Although we found similar patterns of neural activity in these neurons, our simultaneous force measurements reveal that VTA GABA neurons do not encode reward prediction. In our experiments, distinct firing patterns were observed depending on the direction of force exertion, while reward prediction remained the same, as measured by anticipatory licking rate. When the spout was placed in front of the mouth, mice exerted forward force, and many VTA GABA neurons were activated (Forward+ and Bidirectional+), while some GABA neurons were inhibited (Backward+ and Bidirectional-). The opposite pattern was observed when backward force was generated in the “spout behind” condition. Importantly, VTA GABA neurons also maintained their force tuning even in the absence of reward-predicting cues and licking behavior, even in spontaneous behavior.

Eshel et al. (2015) argue that the activation of VTA GABA neurons, which reduces DA neuron activity, might update the cue value and the reward prediction error. However, when we stimulated VTA GABA neurons, there was a robust suppression in the performance of both anticipatory and consummatory behaviors, including force exertion and licking. This effect did not influence the learned association, as mice showed immediate recovery in performance on subsequent trials in the absence of stimulation (Figure 6 and Figure S9).

Some studies found that VTA GABA neurons are also excited by aversive stimuli like foot shocks (Root et al., 2020; Tan et al., 2012), air puffs (Cohen et al., 2012), and aerial threat-related visual stimuli (Z. Zhou et al., 2019). These studies, however, did not measure force exertion or kinematics, the equivalent of force in freely moving animals. We also delivered air puffs to the face while measuring force exertion. On air puff trials, the force tuning of VTA GABA neurons remains similar, though the behavior is different. In response to air puffs, mice showed immediate backward force, followed by a forward rebound after the air puff, which was neglected in previous studies. VTA GABA neurons maintain their force-tuning properties in backward retreat behavior and forward rebound to air puffs. It is worth noting that Cohen et al. also showed similar diversity in Type II and Type III neurons’ response to air puffs, as shown in their Figure S9 (Cohen et al., 2012), but lacking force measures they were not able to interpret this pattern of neural activity.

Studies have suggested that optogenetic stimulation of VTA GABA neurons can induce place aversion (Tan et al., 2012) and suppress consummatory behaviors (van Zessen et al., 2012). These studies used various stimulation protocols, ranging from 4 Hz (Lowes et al., 2021), 20 Hz (W.-L. Zhou et al., 2022), 40 Hz (Eshel et al., 2015), and 60 Hz (Z. Zhou et al., 2019) to continuous laser stimulation (Tan et al., 2012; van Zessen et al., 2012), to activate GABA neurons. Other studies found that chemogenetic activation of these neurons inhibits instrumental response to reward-predictive cues (Wakabayashi et al., 2019). Here we did not observe clear aversion effects. Whether this is due to different stimulation parameters or experimental setups remains to be determined.

Van Zessen et al. (2012) did not observe reduced anticipatory licking when they stimulated VTA GABA neurons during the CS-US interval; they only observed interruption of reward consumption when stimulating after US delivery. They concluded that VTA GABA activation selectively reduces consummatory behavior but not anticipatory behavior. However, we found that this suppression is not specific to reward consumption. When we eliminated consummatory behavior by omitting rewards, stimulation still suppressed force exertion (Figure 7). We therefore conclude that the net effect of VTA GABA activation is a general suppression of force exertion rather than a selective suppression of consummatory behavior. The discrepancy between their study and ours could be explained by their use of a long delay (5s) between CS and US. It is well-known that, in delay conditioning procedures, the latency of CR increases as the delay increases (inhibition of delay) (Pavlov, 2010). This is true in Van Zessen et al. (2012), as their mice showed anticipatory licking (CR) usually towards the end of the CS. Thus effects on anticipatory response could be obscured by the long delay. When a shorter delay (1.5 s from CS onset, including a 0.5 s trace interval) was used by Eshel et al. (2015), the anticipatory licking was significantly suppressed by the excitation of VTA GABA neurons, similar to what we observed.

The stimulation effects also agree with observed effects in freely moving mice. Without constraints, force generation is expected to be translated into movement kinematics. Hughes et al. (2019) found optogenetic stimulation could reliably cause head movements in freely moving mice, which reduced licking. As technical limitations did not allow us to measure rotational movements and associated torques, we cannot rule out the possibility that VTA GABA neurons can also generate torque responsible for rotational kinematics.

### Limitations of the study

In our experiments, optogenetic excitation should activate all VTA GABA neuron subpopulations and the behavioral effects should be the net results of exciting all GABA neurons. However, it is unclear why a net backward force was generated by stimulation. It is possible that some populations (e.g. Backward+) have a much greater influence on downstream circuits and effectors than other populations. As the detailed connectivity of these functionally defined neuronal populations remains unknown, it is difficult to determine the causal role of each population in generating motor output.

It should also be noted that force generation quantified here cannot be equated with torque generation by specific muscles. Like kinematic variables, the type of force representation observed here is a high-level command that is sent to lower levels to generate specific motor outputs. It could involve many muscle groups. As VTA GABA neurons are known to project to many different areas, future work will be needed to determine how such a top-down command signal ultimately generates movement.

## Acknowledgments

This research was supported by NIH grant NS094754 and DA040791.

## Author contributions

QJ, RNH, and HHY designed the experiments. QJ, RNH, and BL performed surgeries, *in vivo* electrophysiological and optogenetic experiments, and histological analysis. QJ, KB, and RNH performed data analysis. QJ, RNH, and HHY wrote the manuscript. All authors have read and approved the manuscript.

## Declaration of Interests

The authors declare no competing interests.

## STAR★Methods

### Key resources table

| REAGENT or RESOURCE | SOURCE | IDENTIFIER |
| --- | --- | --- |
| Bacterial and virus strains |  |  |
| pAAV5-EF1a-DIO-hChR2(E123T/T159C)-eYFP | Addgene | #35509 |
| pAAV1_hSyn1-SIO-stGtACR2-FusionRed | Addgene | #105677 |
| pAAV-FLEX-GFP | Addgene | #28304 |
| Experimental models: Organisms/strains |  |  |
| Mouse: VGAT-ires-cre | Jackson Labs | Mouse Strain: 016962 |
| Mouse: DAT-ires-Cre: 1tm2(cre)Lowl/J | Jackson Labs | Mouse Strain: 006660 |
| Mouse: Ai32: B6;129S-Gt(ROSA)26Sortm32(CAGCOP4*H134R/EYFP)Hze/J | Jackson Labs | Mouse Strain: 012569 |
| Software and algorithms |  |  |
| MATLAB 2022b | MathWorks | https://www.mathworks.com/products/matlab.html |
| GraphPad Prism 9 | GraphPad | https://www.graphpad.com/scientific-software/prism/ |
| Offline Sorter 3.0 | Plexon | https://plexon.com/products/offline-sorter/ |
| NeuroExplorer 5.414 | Nex Technologies | http://www.neuroexplorer.com/downloadspage/ |
| Cerebus Central Suite - Fiber Optic | Blackrock Neurotech | https://blackrockneurotech.com/research/support/software/ |
| Other |  |  |
| DAPI Fluoromount-G | Southern Biotech | Cat# 0100-20 |

### Animals

All experimental procedures and protocols were approved by the Animal Care and Use Committee at Duke University (Protocol # 162-22-09). 18 VGAT-ires-cre mice, 4 DAT-Cre + Ai32 were used (Jackson Labs, Bar Harbor, ME). DAT::Ai32 mice were generated by crossing Ai32 mice, which express channelrhodopsin (ChR2) in neurons with Cre, and DAT-Cre mice, which express Cre under the dopamine transporter (DAT) promoter (Madisen et al., 2012). Mice (2-10 months old) were housed on a 12:12 light cycle. Both males and females were used (13 males and 9 females). Because no significant sex difference was found, we combined data from male and female mice. Mice implanted with optrodes were singly housed, while mice implanted with optic fiber were group-housed. Experiments were conducted during the light phase. During experiments, mice were placed on water restriction and maintained at 85% of their initial body weights. Mice received free access to water for approximately two hours each day. Surgeries, experiments, and access to water were conducted during the light phase.

## Method details

### Viral Construct

pAAV5-EF1a-DIO-hChR2(E123T/T159C)-eYFP(Addgene plasmid, #35509), pAAV1_hSyn1-SIO-stGtACR2-FusionRed(Addgene viral prep, #105677), and pAAV-FLEX-GFP(Addgene plasmid, #28304) were used in this study.

### Stereotactic surgeries

Mice were anesthetized with 2.0 -3.0% isoflurane, received Meloxicam (2mg/kg) via intraperitoneal injection, and placed into a stereotactic frame (David Kopf Instruments, Tujunga, CA). They were maintained during surgery at 1.0-1.5 % isoflurane with oxygen flowing at 0.6L/min for the duration of the surgery. A craniotomy was then drilled above the VTA: anterior-posterior (AP): −3.2 to –3.4 mm, medial-lateral (ML): ±0.3 to 0.6 mm relative to bregma). 200-400 nl of pAAV5-EF1a-DIO-hChR2(E123T/T159C)-eYFP or pAAV1_hSyn1-SIO-stGtACR2-FusionRed or pAAV1-FLEX-GFP were unilaterally injected into the VTA for in vivo electrophysiology experiments, or bilaterally for optogenetic stimulation experiments (AP: −3.1 to –3.4 mm, ML: ± 0.3 to 0.6 mm, dorsal-ventral (DV): −4.0 to −4.6 mm relative to bregma) at a rate of 1 nl/s (Nanoject 3000, Drummond Scientific). Once the target coordinates were reached, the pipette was left to sit for 10 minutes at the injection site to allow absorption of the virus. 2–3 drops of Bupivicane were applied to the wound.

For electrophysiological experiments, an optic fiber was attached to the electrode array at an angle (∼15°). The electrodes were slowly lowered onto the VTA (AP: −3.1 to −3.4 mm, ML: ±0.3 to 0.6 mm, DV: − 4.1 mm relative to bregma) and grounded to four cranial screws by a silver grounding wire. For optogenetic stimulation experiments, two custom-made optic fibers (5 − 6 mm length below ferrule, > 70% transmittance, 105 μm core diameter) were then implanted at an angle (15°) above the VTA (AP: −3.1 to −3.4 mm, ML: ±1.6 mm, DV: −3.8 to 4.1 mm). Fibers and electrodes were secured to the skull using screws and dental acrylic, and all mice were fitted with a steel head implant for head fixation. All mice were allowed to recover for two weeks before beginning behavioral experiments.

### Histology

Mice were transcardially perfused with 0.1M phosphate buffered saline (PBS) followed by 4% paraformaldehyde (PFA) in order to confirm viral expression as well as optic fiber and electrode placement. To confirm placement, brains were post-fixed in 4% PFA for 72 hrs. Tissue was stored for 48 hours in 30% sucrose for cryoprotection before cryostat sectioning (Leica CM1850) at 60 µm. Sections were mounted and immediately coverslipped with Fluoromount G with DAPI medium (Electron Microscopy Sciences; catalog no. 17984-24). To validate placement of electrodes and fibers, fluorescent images were acquired and stitched using AXIO Zoom V16 (Zeiss).

### Head-fixed behavioral system

We developed a customized head-fixation device for measuring forces exerted by the mice during behavioral testing and stimulation, as previously described (Hughes, Bakhurin, Barter, et al., 2020). The head was clamped via a bar implanted into the dental cement onto the head-fixation frame, which contained 3 load cells (RB-Phil-203, RobotShop.com). The mice were positioned on an elevated, enclosed perch that also contained 2 load cells. These load cells detected downward forces exerted by either the left or right feet, but data from foot forces were not included in this study. Load cells measure force by converting mechanical deflections to voltage signal, which is sampled at 1 kHz and amplified using an INA125P (Texas Instruments). Each load cell was arranged orthogonally to detect forces exerted in the up-down, left-right, and forwards-backward directions. Load cell voltages, electrophysiological data, and timestamps for licks, reward, and photo-stimulation were recorded with a Blackrock Cerebus system (Blackrock Microsystems) for offline analysis.

For delivery of sucrose rewards, a spout connected to a reservoir with a 10 % sucrose solution was positioned close to the mouth. Reward delivery was controlled by opening a solenoid valve (161T010, NResearch, NJ) attached to the tubing connected to the spout. A capacitance-touch sensor (MPR121, AdaFruit.com) attached to the spout was used to detect licks. A 3 kHz buzzer emitting an 80dB sound was stabilized approximately 35 cm in front and 20 cm above the head-fixation frame, on the wall of the behavioral box.

### Stimulus-reward trace-Conditioning Task

Mice were first allowed to habituate to the head-fixed condition and trained to receive water rewards delivered manually by the experimenters for 5 minutes for approximately 1-2 days. Once mice were habituated to head fixation, a water spout containing 10 % sucrose was placed underneath their nose at in front and behind locations (±1 mm change in position). In order to eliminate the confounding variable of the sound of the solenoid during reward delivery, 50 dB white noise was continually present in the background. The mice were trained for 70-90 trials/day on a Pavlovian trace conditioning task for 10-14 days before electrophysiological recording or optogenetic stimulation experiments. At the beginning of each trial, a 3 KHz tone that lasted 200 ms was presented, followed by delivery of 10-20 ul of 10% sucrose. The delay between the onset of the tone and reward delivery was 1s. There was a random intertrial interval that varied between 7 and 60 seconds. For the sessions with reward omissions, trials have a 30-50% probability of omission. Experiments using aversive air puffs were conducted in separate sessions. Air puff was delivered using an EFD 1500 XL pneumatic fluid dispenser. The output tube was placed 20 mm in front of the face, and the air puff lasted 20 ms.

### Wireless *in Vivo* electrophysiology

Drivable electrodes were single-drive movable micro-bundles of tungsten electrodes (1 x 16; 23 μm diameter) placed within a guide cannula (Innovative Neurophysiology, Inc.). Electrophysiological data were recorded using a miniaturized wireless head stage (Triangle Biosystems) that was interfaced with a Blackrock Cerebus data acquisition system (Blackrock Microsystems). A digital bandpass filter was applied to the electrophysiological data (250 Hz – 5 kHz), and spike timestamps and waveforms were recorded at 30 kHz. Filtered data were sorted using Offline Sorter (Plexon). A 3:1 signal-to-noise ratio and an 800 μs or greater refractory period were required for the neural data to be used for analysis. Single units were selected based on a principal component analysis of waveforms using 2 principal components. To acquire new neurons with drivable electrodes, the electrodes were lowered by 25 μm at a rate of 1 μm per second, repeated 2-3 times. Peri-event raster plots were generated using NeuroExplorer (Nex Technologies).

### Optogenetic experiments

Optogenetic stimulation sessions were identical to the Pavlovian conditioning tasks with electrophysiological recordings as described above. Pulses of light (Excitation: 5-8 mW measured at the tip of the optic fiber connected to the optic implant, 15 ms pulse width, 40 Hz 40 pulses) were delivered via a laser (470 nm DPSS laser, Shanghai Laser & Optics) and controlled through a custom MATLAB script during the behavioral session. In Pavlovian conditioning stimulation experiments (80-100 trials per session), every trial was rewarded, and stimulation was delivered with a 20% probability at CS or US, respectively. In reward omission with stimulation experiments (80-100 trials per session), each trial has a 50% probability of reward delivery and a 50% probability of stimulation at 1 second after CS. Probability was controlled pseudo-randomly by MATLAB.

### Quantification and statistical analysis

All analyses were performed with either MATLAB 2023a, NeuroExplorer, or Graphpad Prism 10. Statistical analyses were performed in MATLAB 2023a or Graphpad Prism 10. The sample size and statistical tests used were indicated in the figure legends. The data are shown as the mean ± standard error of the mean (SEM). ∗p < 0.05; ∗∗p < 0.01; ∗∗∗p < 0.001; ∗∗∗∗p < 0.0001; ns, nonsignificant (p > 0.05).

### Force conversions

Detailed instructions for the setup of load cell systems and conversion of signals can be found in previous publications (Bakhurin et al., 2020; Hughes, Bakhurin, Barter, et al., 2020). Briefly, we calibrated the load cell circuits using a conversion factor (expressed in Newtons per Volt) determined by the linear relationship between the voltage changes resulting from known masses placed on the sensor. Force was determined by multiplying the voltage signal by the conversion factor to obtain a value in Newtons.

### Detection of movements

Forward and backward movement start times and end times were identified by changes in force exertion, surpassing a threshold of three standard deviations, a duration longer than 100 ms, and at least 100 ms from another movement. Licking was detected by the contact of spout by the capacitance-touch sensor.

### Functional classification of VTA neurons

Previous studies have shown that VTA dopamine and GABA neurons have distinct firing patterns (Cohen et al., 2012; Morales & Margolis, 2017). For each recorded neuron, the firing rate was calculated within the window of 1 second before the CS and 2 seconds after the CS, using 10 ms bins. The baseline of each unit during trials of different spout placements was determined by the average firing rate of the first 50 bins of each trial. The firing rates of all neurons were baseline-subtracted, z-scored, and combined into a matrix for the front and behind spout placements. We then applied agglomerative clustering to the functional vector matrix of all cells, resulting in five distinct response profiles. All preprocessing was conducted in Matlab, and clustering was performed using the ‘clusterdata’ function with Euclidean distance and Ward linkage parameters. After clustering, we manually verified the consistency of the responses.

### Electrophysiological and optogenetic identification of VTA GABA neurons

The firing activities of classified groups resembled those of neuron types reported in previous studies, with dopamine neurons showing bursting activities at CS and US and GABA neurons maintaining either elevated or reduced activities during CS-US interval, depending on spout locations. We examined the electrophysiological properties of 5 clusters. The electrophysiological properties of putative GABA and Unclassified neurons agreed with the respective characteristics reported in previous studies *in vivo* (Bouarab et al., 2019; Morales & Margolis, 2017).

Optogenetic tagging was further used to confirm GABA neuron identity. Light (5-8 mW, 5-15 ms pulse width, 10-40 Hz for 1 second for ChR2, 500-1000 ms pulse for inhibition) was delivered via laser (470 nm DPSS laser, Shanghai Laser & Optics) before or after the behavioral session of VGAT-Cre mice. For mice injected with ChR2, we examined spikes within 10 ms of the onset of light pulses. Neurons were classified as tagged if the first pulses of light produced a spike that occurred within 7 ms of stimulation, evoked waveforms highly correlated with spontaneous waveforms (R2 ≥ 0.95), and >50% fidelity to firing an action potential following the first pulses. For mice injected with stGtACR, we compared the firing rate during laser and baseline firing rate. Neurons were considered tagged if the firing rate during stimulation showed a decrease of at least 30% compared to the baseline firing rate.

### Analyses of force tuning

Neural and behavioral data from each session were binned (10ms). The peri-event interval was -1 to +2 seconds around CS and spontaneous movement start times or -0.5 to +1 seconds around air puff delivery times. For Forward+ and Backward+ neurons, forward and backward forces were analyzed together. For Bidirectional+ and Bidirectional-neurons, the absolute value of forward and backward forces was analyzed. A cross-correlation was computed between the normalized firing rate and force to determine the time shift between the two signals across the session by the *xcorr* function in MATLAB, using force as the reference. We thus determined the lag of the maximum value of the cross-correlation for positively correlated neurons and the lag to the minimum value for anti-correlated neurons. Neural activity was then shifted according to the lag value. We next sorted the firing rate and excluded outliers (2.5th and 97.5th percentile), which contain very few replicates. We created 20 equal bins with normalized firing rate bins (from -1 to 1s for Forward+ and Backward+ neurons; 0 to 1s for Bidirectional+ and Bidirectional-neurons). Force measures obtained concurrently with the firing rate from each bin were averaged. The non-linear regression (Sigmoidal) was calculated between binned neural activity and the mean force signal of each bin. Force tuning during spontaneous movements and air puffs were analyzed similarly.

### Analyses of behavior in optogenetic experiments

Force signals and lick timestamps were aligned to the onset of CS in different stimulation conditions as described in the main text. The peri-event interval was -1 to +4 seconds around CS (10ms bin). The average force signal and licking behavior during the CR and UR were calculated by averaging across all trials and subtracting the baseline, defined as the mean during the 1 second preceding the CS onset. This resulted in one value for each animal’s licking or force for each optogenetic stimulation condition. Paired t-test was used for the comparison between the two different trial types. One-way ANOVA was used to compare three different trial types. All tests were two-tailed, and a p-value less than 0.05 was considered statistically significant.

## Notes

### Competing Interest Statement

The authors have declared no competing interest.

