## Supplementary figures for "GABAergic neurons from the ventral tegmental area represent and regulate force vectors"

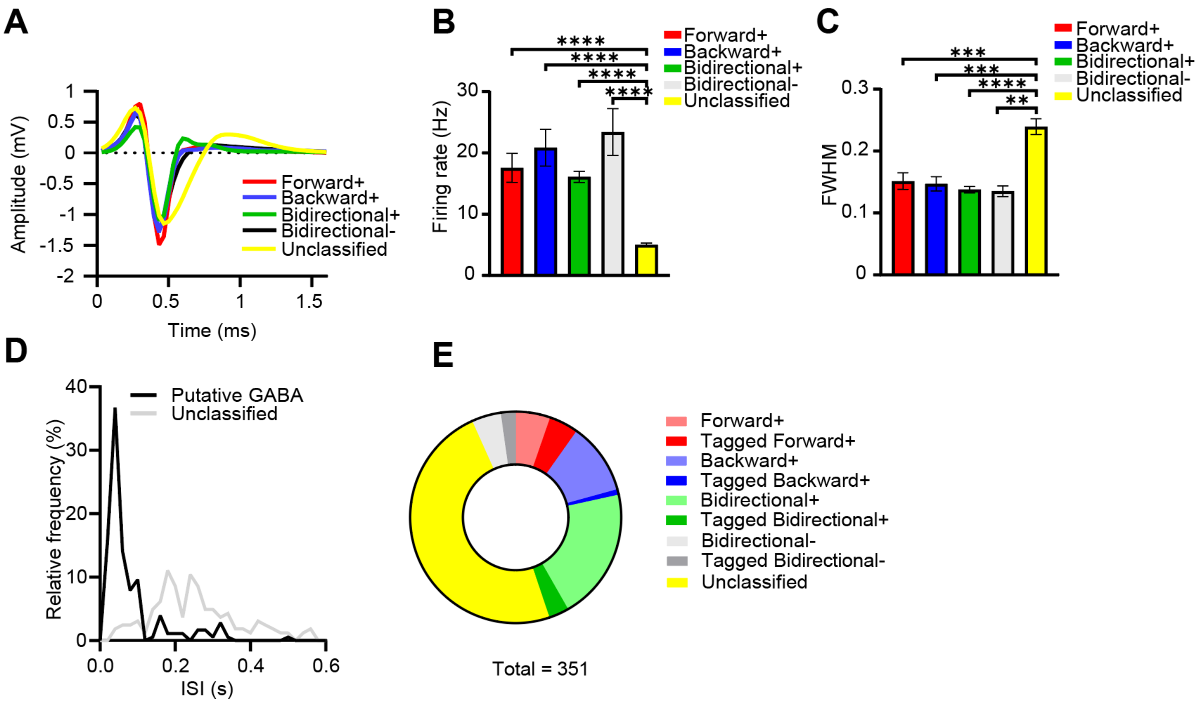


**Figure S1. Electrophysiological properties of VTA neurons.**

**(A)** Average waveforms for different types of putative GABA neurons and unclassified neurons. The unclassified neurons are presumably a mixture of putative dopaminergic and glutamatergic neurons.

**(B)** The bar plot represents the mean baseline firing rates (± SEM) of each neuron type. A one-way ANOVA revealed a significant difference in baseline firing rates across the groups ($F (4, 346) = 41.04$, $p<0.0001$). Post-hoc analyses indicated that the firing rates of all four GABA neuron groups were significantly higher than that of the Unclassified group. **** indicates $p < 0.0001$ compared to the Unclassified group.

**(C)** Full width at half maximum (FWHM; mean ± SEM) for different types of putative GABA neurons and unclassified neurons. A one-way ANOVA revealed a significant difference in FWHM across the groups ($F (4, 346) = 13.16$, $p < 0.0001$). Post-hoc analyses indicated that the FWHM of all four GABA neuron groups was significantly smaller than that of the Unclassified group. The specific p-values for each comparison are as follows: Forward+ vs. Unclassified ($p = 0.0008$), Backward+ vs. Unclassified ($p = 0.0001$), Bidirectional+ vs. Unclassified ($p < 0.0001$), Bidirectional- vs. Unclassified ($p = 0.0027$).

**(D)** Distribution of inter-spike interval (ISI) of GABA neurons and Unclassified neurons.

**(E)** Portions of different types of putative GABA neurons and Unclassified neurons

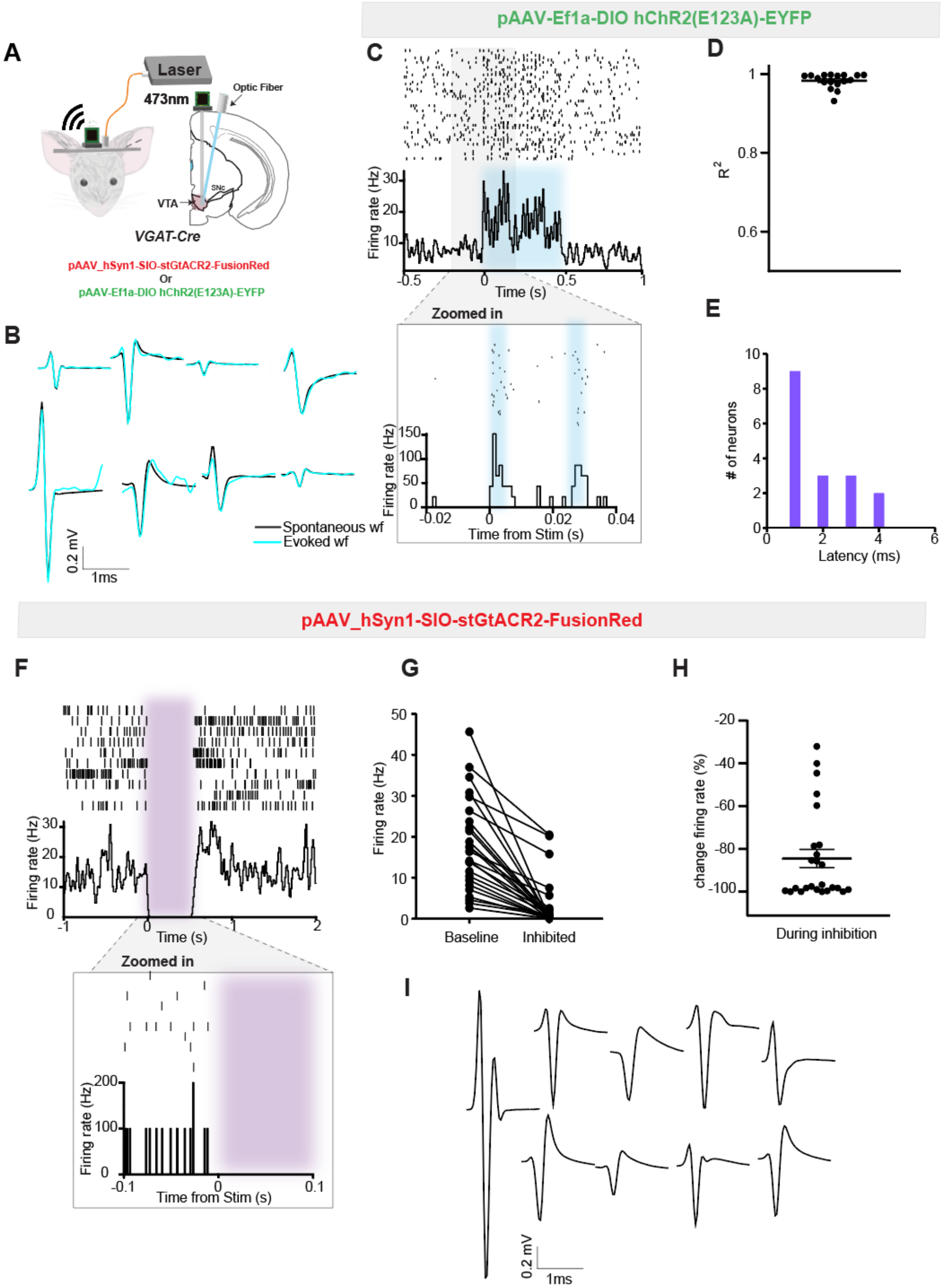


**Figure S2. Opto-tagging using both optogenetic inhibition (stGtACR2) and excitation (ChR2) identified VTA GABAergic neurons.**

**(A)** An optrode was implanted into the VTA of VGAT-Cre mice, and either DIO-stGtACR2 or DIO-ChR2 were injected to tag VTA GABA neurons.

**(B)** Representative ChR2-tagged neurons showing identical spontaneous (black) and light-evoked (cyan) waveforms.

**(C)** Zoomed-in view of a representative tagged GABA neuron using ChR2. Each tick is a spike.

**(D)** Light-evoked waveforms highly correlated with spontaneous waveforms. Each dot is a neuron's correlation value.

**(E)** Latency and number of neurons tagged. All tagged neurons showed modulated within 4 milliseconds (n = 17).

**(F)** Zoomed-in view of a stGtACR2-tagged neuron's response to stimulation.

**(G)** stGtACR2-tagged neurons showed a significant decrease in firing rate compared to baseline (n = 24). Baseline is calculated by the average firing rate during 1 second before the onset of stimulation. Inhibited firing rate is calculated by the average firing rate during stimulation.

**(H)** Change in firing rate is calculated by percentage change from baseline firing rate. Each dot represents a value from a neuron.

**(I)** Example spontaneous wave forms of stGtACR2-tagged neurons.

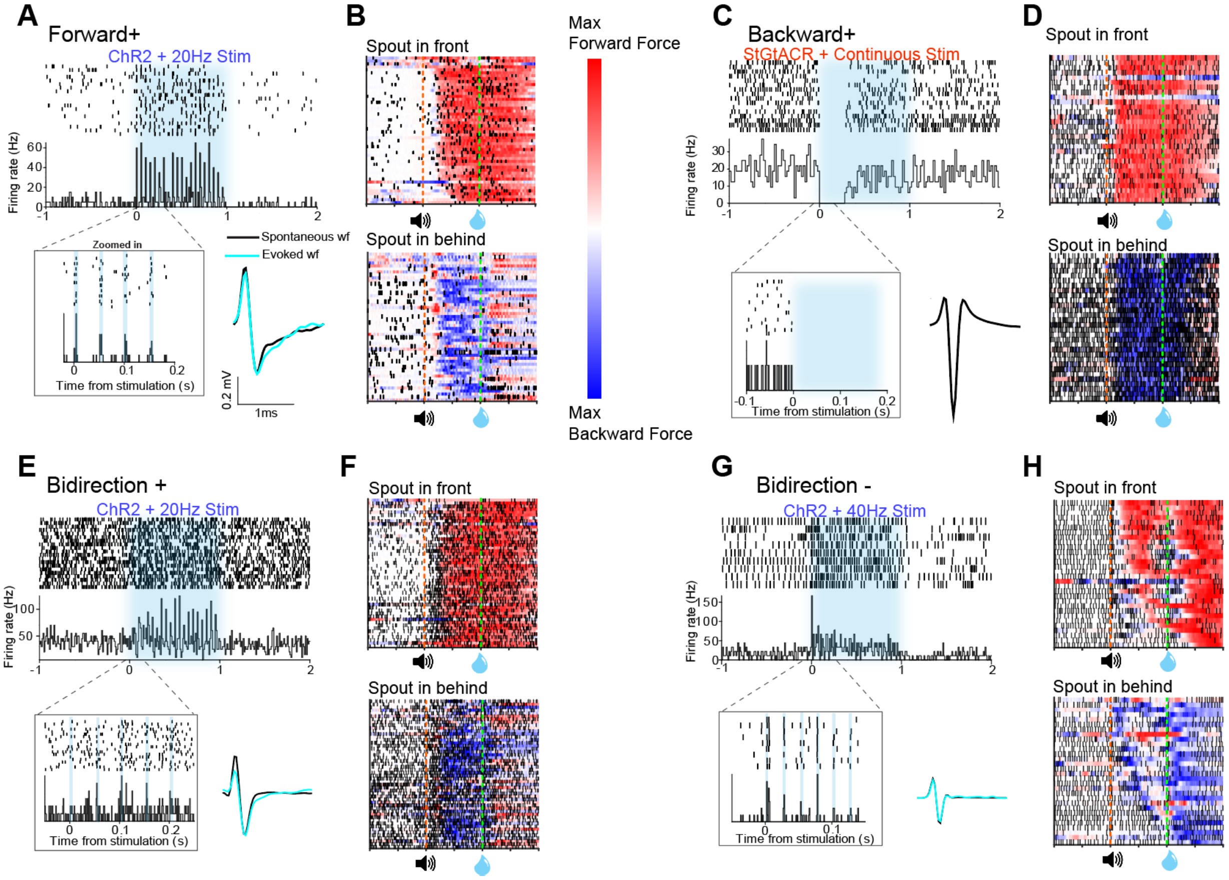


**Figure S3. Example of tagged GABA neuron response.**

**(A)** Response of a Forward+ neuron to stimulation with zoomed-in view; evoked waveforms (cyan) are shown with spontaneous waveforms (black). Each tick is a spike.

**(B)** Spikes and force exertion in trials with different spout location.

**(C-H)** Same tradition as **A-B**, for Backward+ (**C, D**), Bidirectional+ (**E, F**), and Bidirectional- neurons (**G, H**).

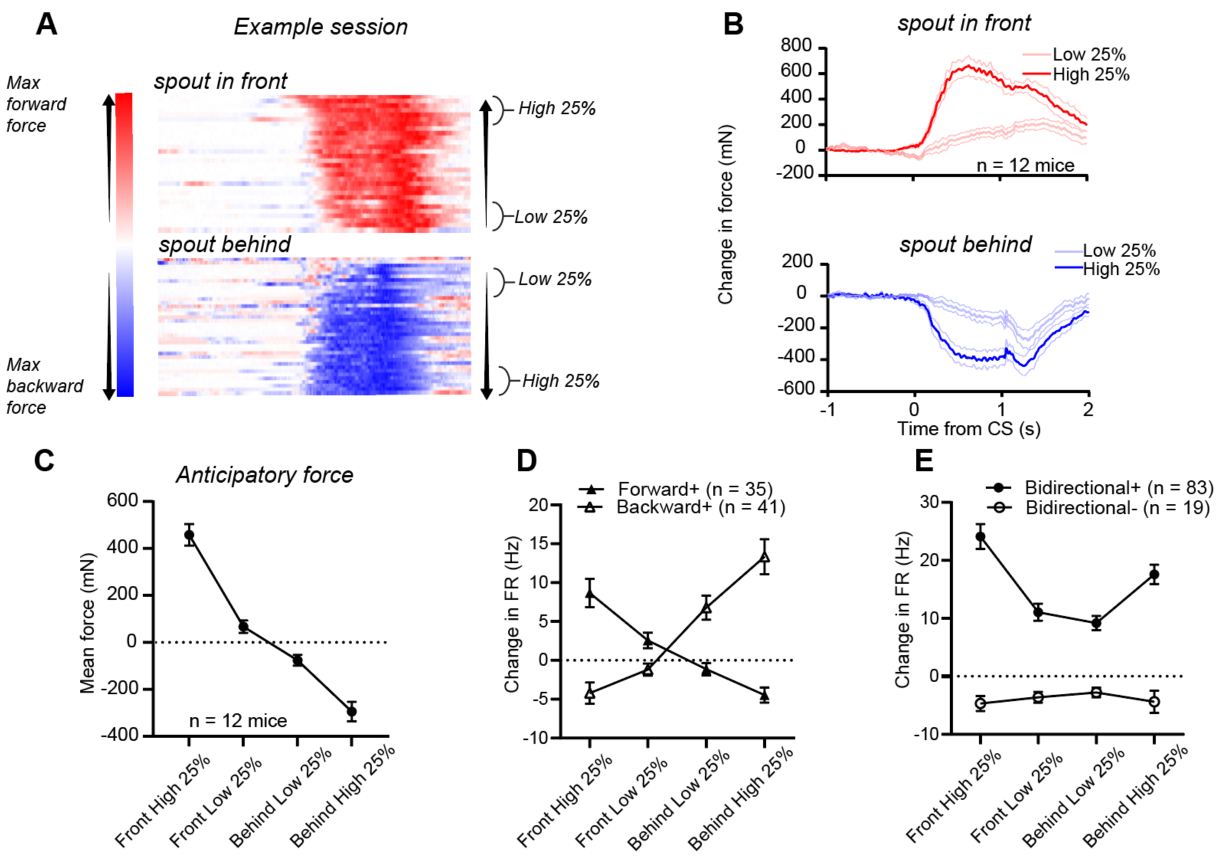


**Figure S4. GABA activities in trials when mice exerted different magnitude of force.**

**(A)** In an example session, the mouse exerted diverse magnitude of anticipatory force throughout the duration of the session. Trials were sorted by anticipatory force magnitude and divided in quartile for each spout placement. Top 25% and bottom 25% of the trials were defined as High 25% and Low 25%.

**(B)** Force exertion during the High 25% and Low 25% trials when the spout was placed in front and behind (n = 12 mice).

**(C)** The average anticipatory force differed across High/Low 25% trials in different spout location conditions. Change in FR is defined by the different between the average firing between CSUS and 1 second before CS.

**(D-E)** Four types of GABA neurons' activities correlated with force magnitude with preference to directions.

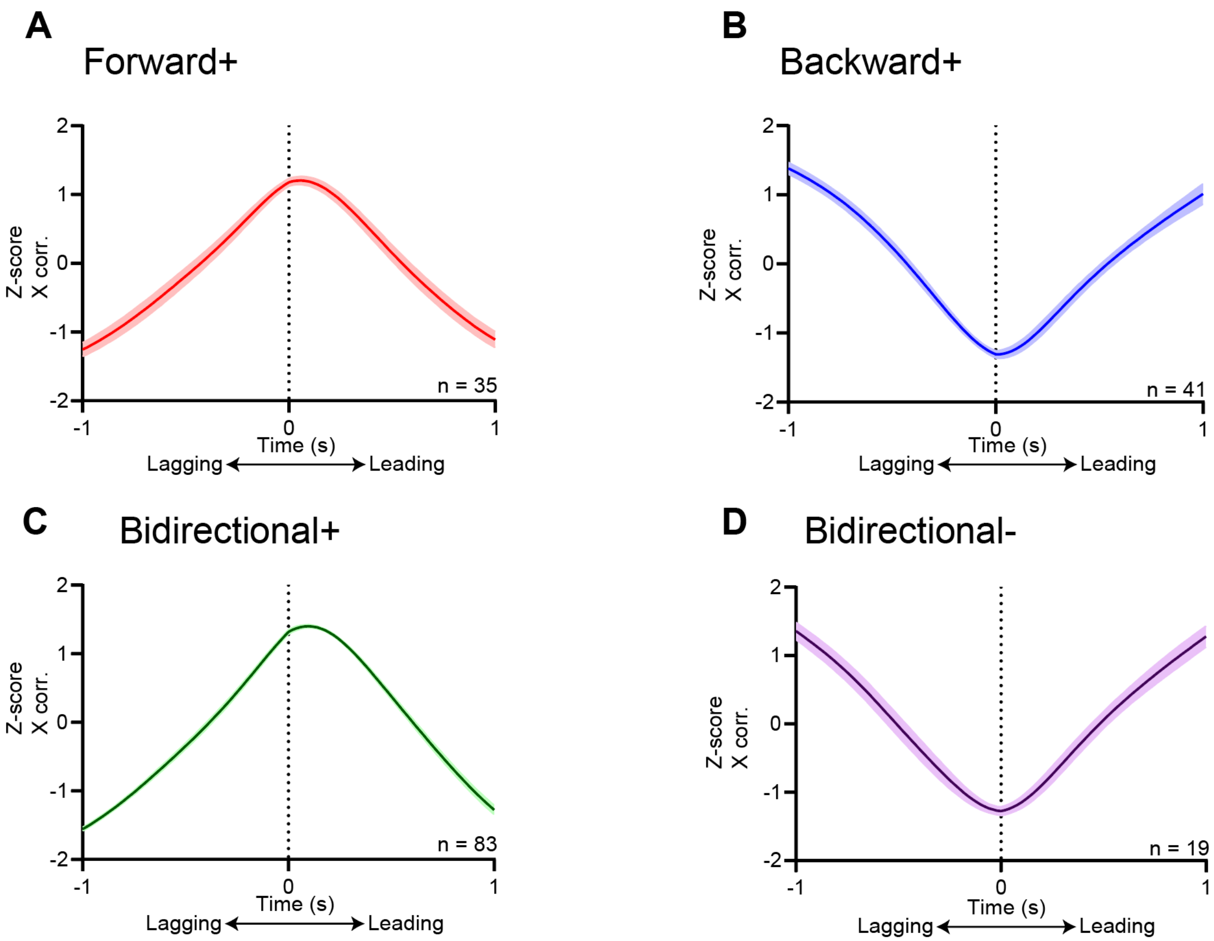


**Figure S5. Temporal relationship between neural activity and force.**
To determine the temporal relationship between firing rate and force, we used cross-correlation using firing rate as the reference variable. Lagging: firing rate changes lag force changes. Leading: firing rate changes lead force changes.
**(A-D)** Forward+ (**A**), Backward+(**B**), and Bidirectional+(**C**) neural activities lead force exertion, while Bidirectional- (**D**) neural activity slightly lags force generation.

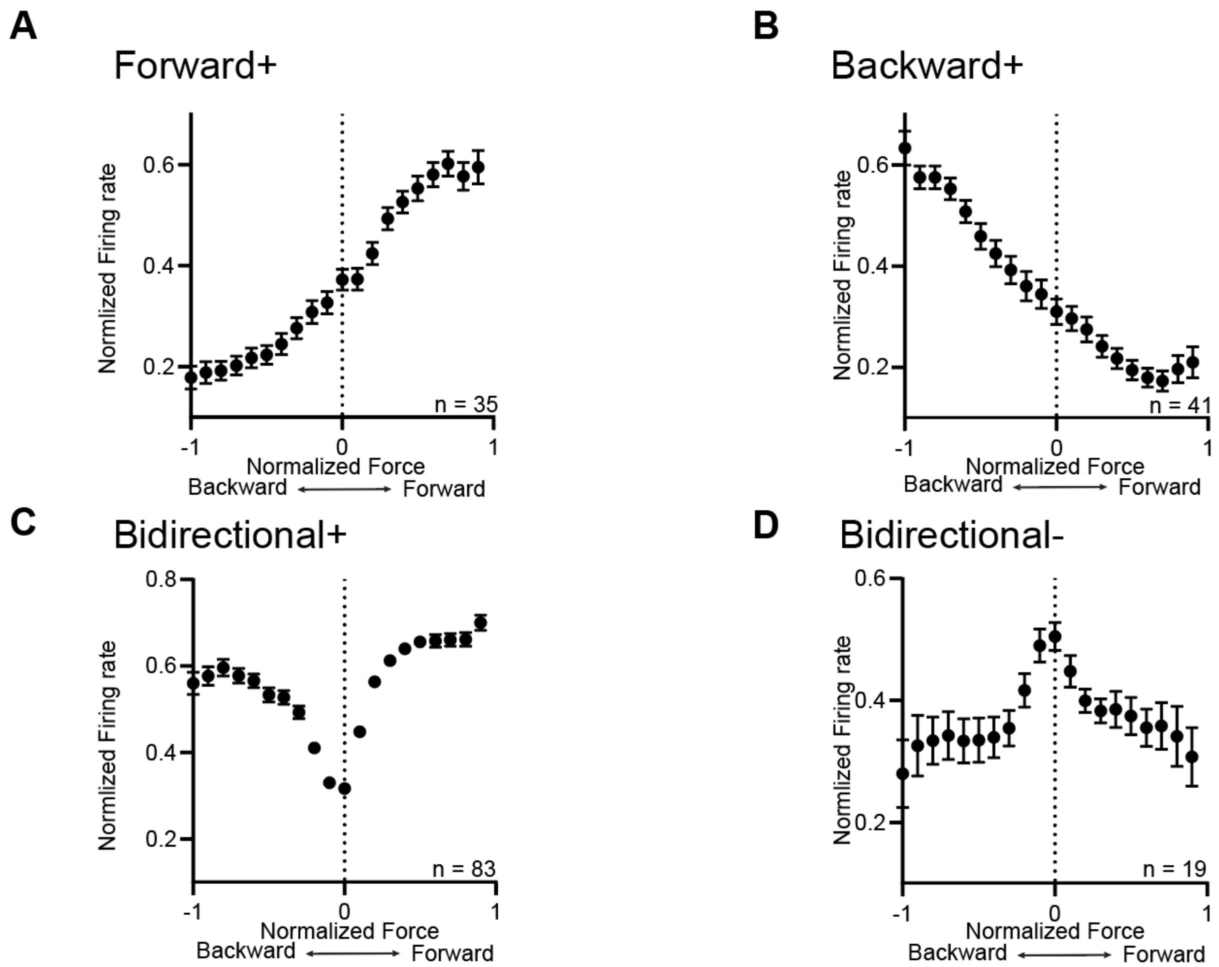


**Figure S6. Different groups of GABA neurons exhibited distinct tuning patterns for force exertion in different directions.**

**(A-D)** In-task backward and forward forces were normalized and binned (-1 for backward max and 1 for forward max values; 20 bins). GABA neurons showed different tuning patterns to force exertions.

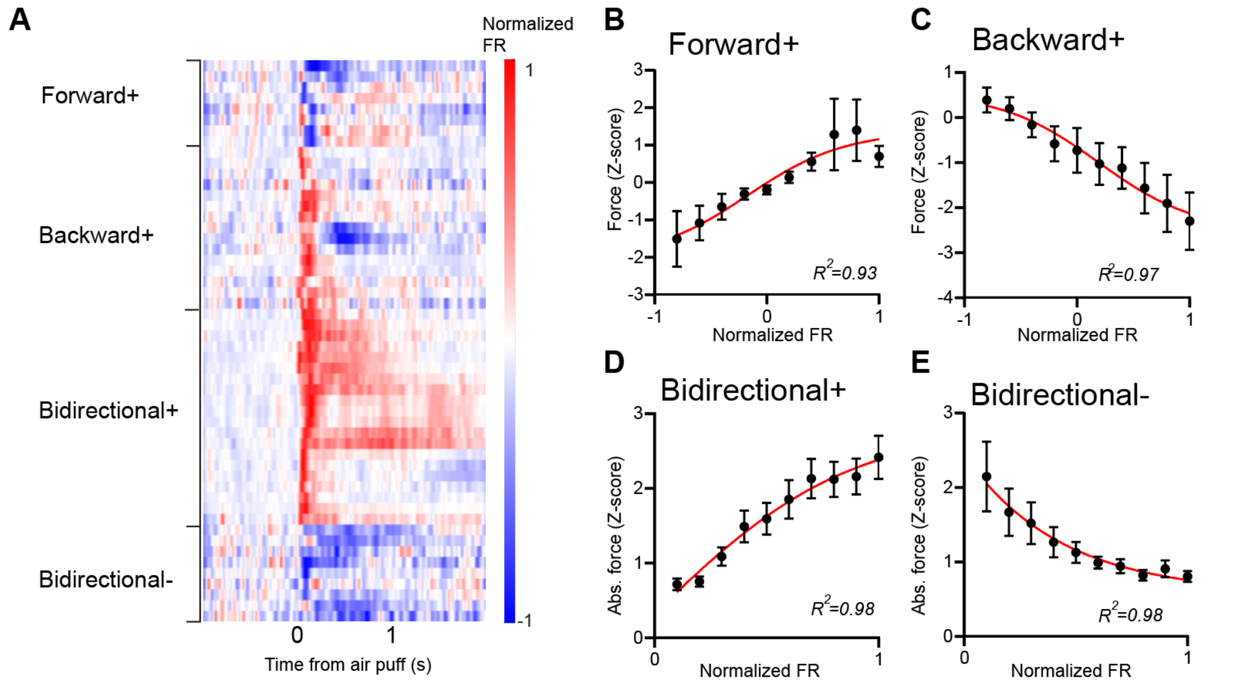


**Figure S7. VTA GABA neurons maintained force tuning in response to aversive air puffs.**

**(A)** Summary of four types of GABA neuron normalized firing rates in response to air puffs.

**(B)** Forward+ population exhibited tuning for forward force ($R^2 = 0.93$, n = 8).

**(C)** Backward+ population exhibited tuning for backward force ($R^2 = 0.97$, n =13).

**(D)** Bidirectional+ population exhibited tuning for absolute force amplitude ($R^2 = 0.98$, n = 22).

**(E)** Bidirectional- population showed a negative correlation with absolute force amplitude ($R^2 =$ 0.98, n = 9).

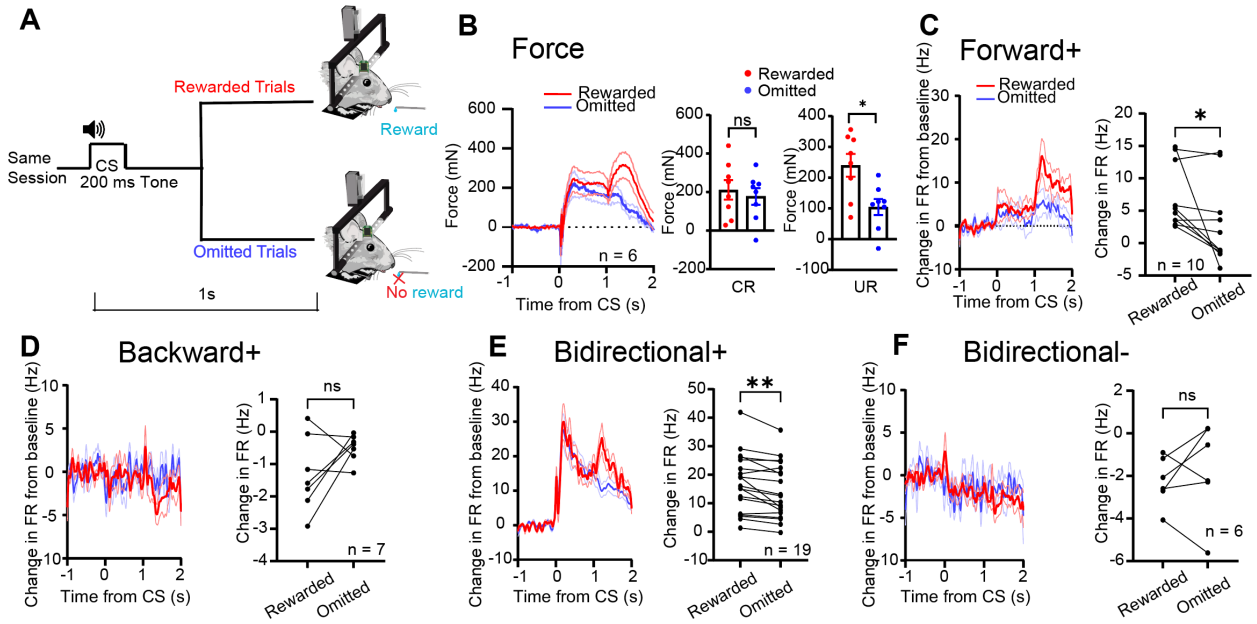


**Figure S8. VTA GABA activity represents force when the reward is omitted.**

**(A)** Schematic illustration of reward omission experiments.

**(B)** *Left*: group summary of force exertion in rewarded trials and omitted trials. *Center*: mice exerted similar force as the conditioned response in rewarded and omitted trials (paired t-test, t = 1.517, p = 0.173, n = 8 sessions). *Right*: mice exerted less force on omitted trials (paired t-test, t = 3.012, p = 0.0196, n = 8 sessions).

**(C)** *Left*: change in firing rate from baseline (average firing rate between -1 to 0s) of Forward+ neurons in rewarded and omitted trials. *Right*: Forward+ neurons had a lower firing rate in omitted trials (paired t-test, t = 2.308, p = 0.0464, n = 10; Change in FR: average FR during UR – baseline).

**(D-F)** Change in firing rate from baseline of Backward+ (paired t-test, t = 1.671, p = 0.146, n = 7), Bidirectional+ (paired t-test, t = 3.812, p = 0.0013, n = 19), and Bidirectional- (paired t-test, t = 0.769, p = 0.677, n = 6) neurons in rewarded and omitted trial.

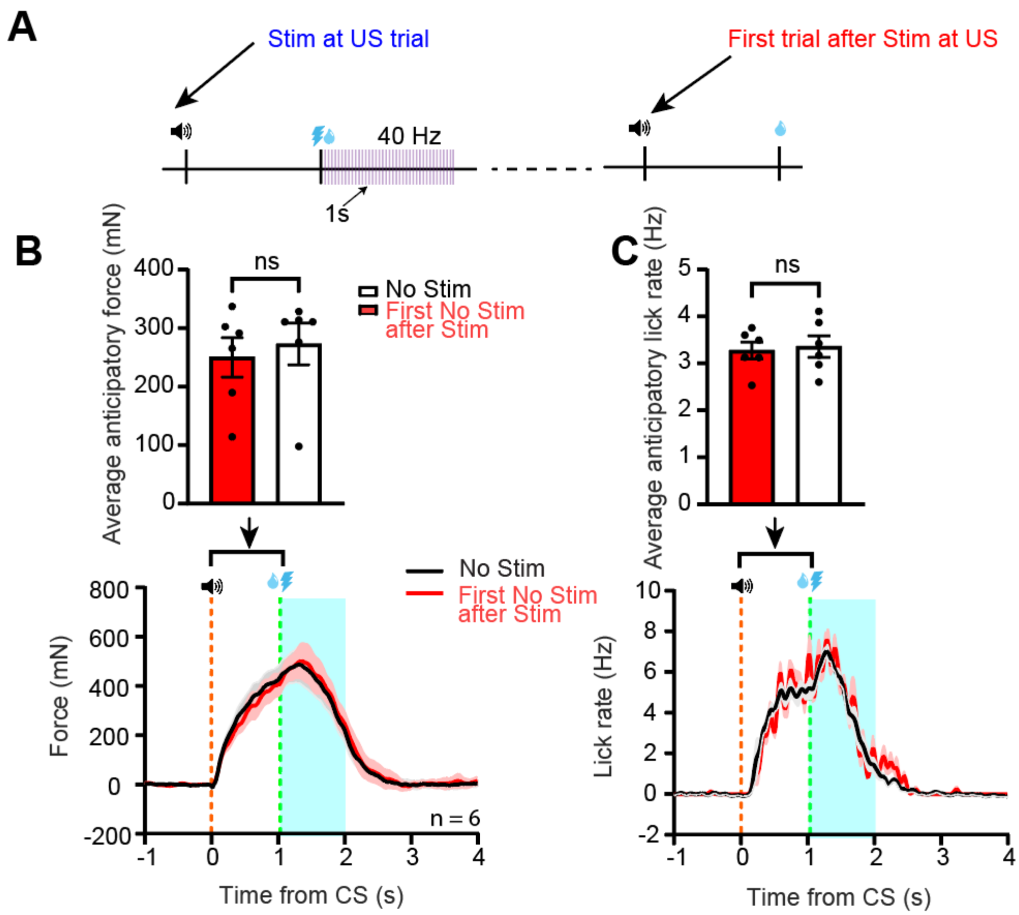


**Figure S9. CR in subsequent trials following stimulation at the reward**.
**(A)** Illustration of the first trial after the trial with stimulation at the US.
**(B-C)** Anticipatory force exertion (paired t-test, $t = 1.458$, $p = 0.2047$, $n = 6$) and licking (paired t-test, $t = 0.5460$, $p = 0.6085$, $n = 6$) were not influenced in the first trial following the trial with stimulation at the US.

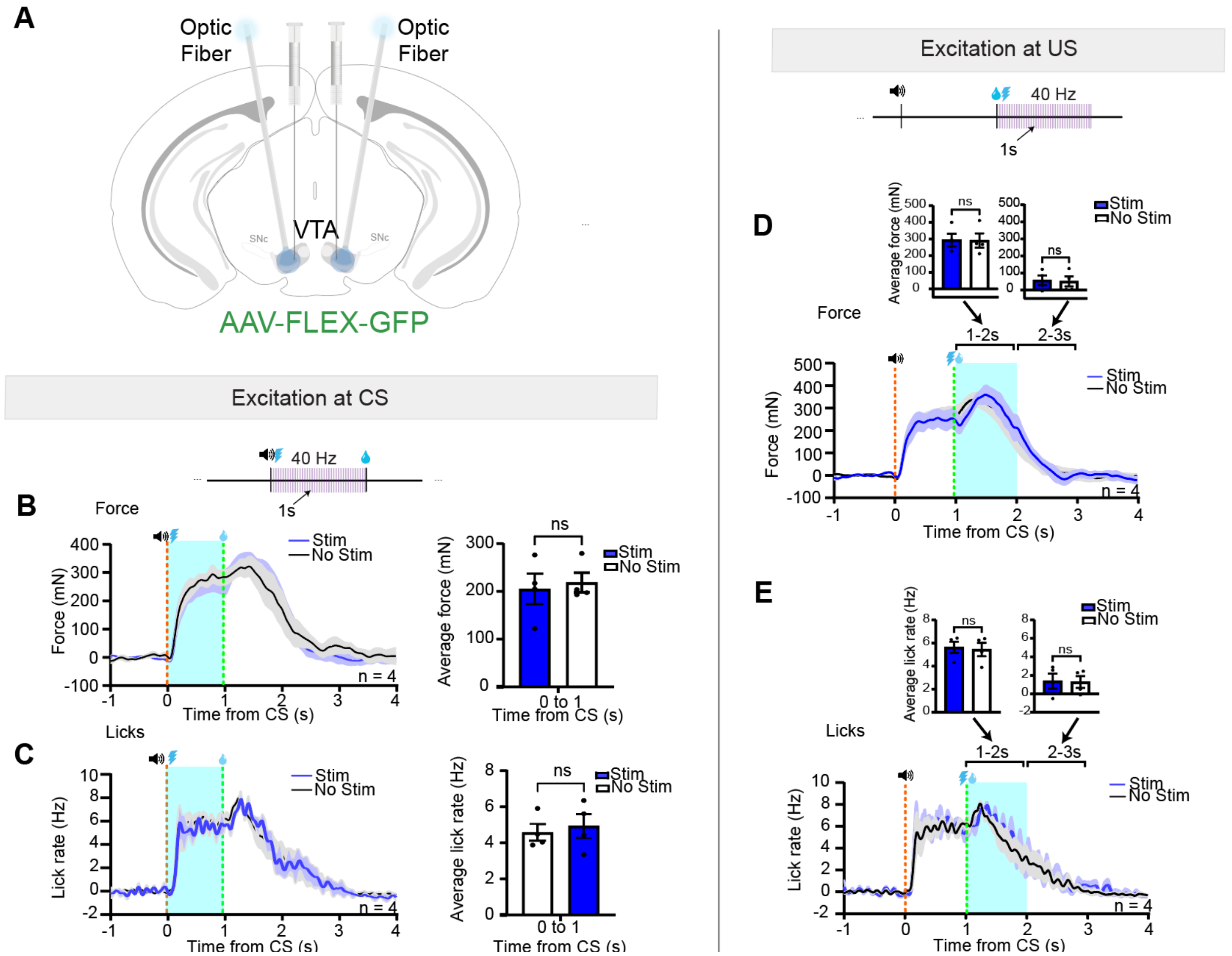


**Figure S10. Control mice showed no behavioral effects during optogenetic stimulation**

**(A)** Schematic illustration of optic fibers implanted into bilateral VTA with AAV-FLEX-GFP injection.

**(B)** Force exertion in control mice was not affected by stimulation delivered at the CS (paired t-test, $t = 0.572$, $p = 0.608$, $n = 4$ mice).

**(C)** Licking rate in control mice was not affected by stimulation delivered at the CS (paired t-test, $t = 1.042$, $p = 0.374$, $n = 4$ mice).

**(D)** Force exertion in control mice was not affected by stimulation delivered at the US (paired t-test, $t = 0.309$, $p = 0.777$, $n = 4$ mice) and after the stimulation (paired t-test, $t = 0.845$, $p = 0.460$, $n = 4$ mice).

**(E)** Licking in control mice was not affected by stimulation delivered at the US (paired t-test, $t=1.771$, $p = 0.175$, $n = 4$ mice) and after the stimulation (paired t-test, $t = 0.537$, $p = 0.628$, $n = 4$ mice).

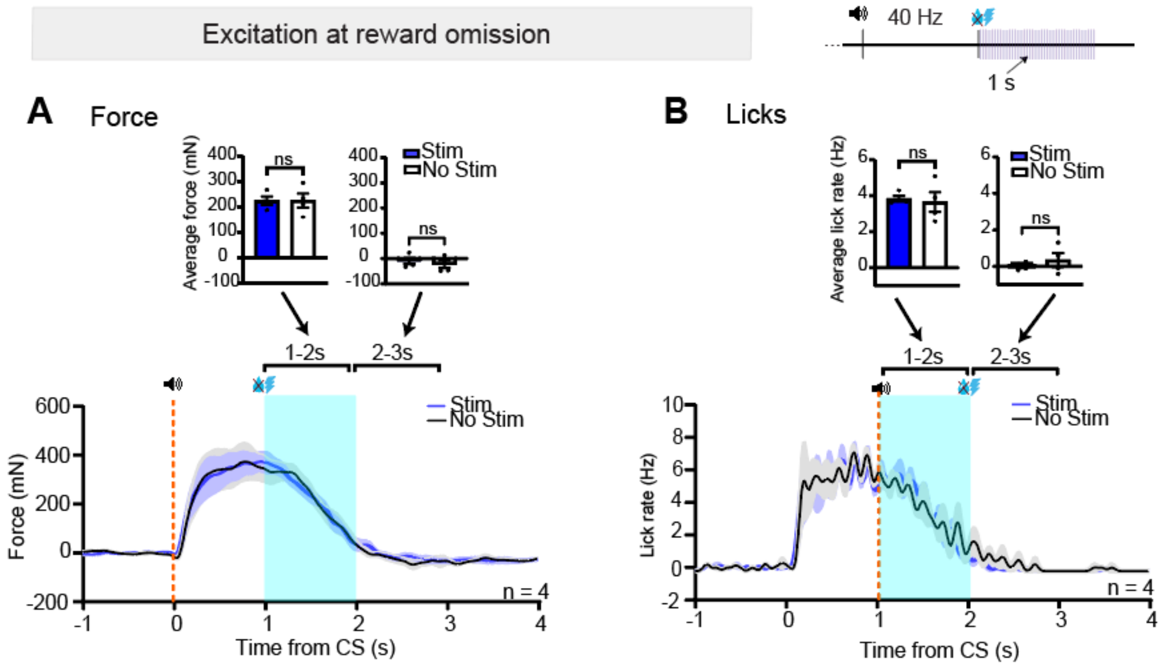


**Figure S11. Control mice showed no behavioral effects during and after optogenetic stimulation-reward omission experiments**

**(A)** Stimulation did not have an effect on force exertion in control mice when the reward was omitted (during stimulation: paired t-test, $t = 0.0288$, $p = 0.979$, $n = 4$ mice; after stimulation: paired t-test, $t = 0.653$, $p = 0.560$, $n = 4$ mice).

**(B)** Stimulation did not have an effect on licking in control mice when the reward was omitted (during stimulation: paired t-test, $t = 0.311$, $p = 0.776$, $n = 4$ mice; after stimulation: paired t-test, $t = 0.967$, $p = 0.404$, $n = 4$ mice).

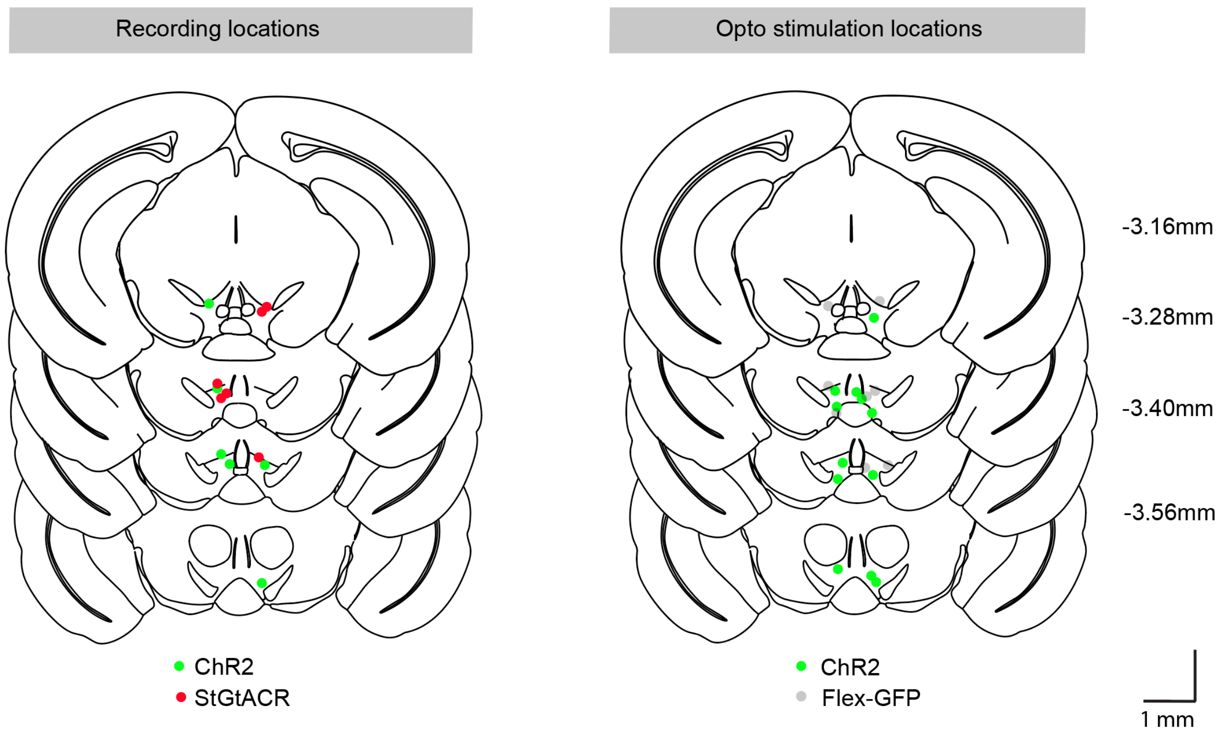


**Figure S12. Verification of stimulation and recording sites in the VTA.**

**Supplementary Video 1**: Optogenetic stimulation of VTA GABA neurons at the tone prevents forward force exertion.

**Supplementary Video 2**: Optogenetic stimulation of VTA GABA neurons after reward delivery suppresses forward force exertion.
